# Aeon: an open-source platform to study the neural basis of ethological behaviours over naturalistic timescales

**DOI:** 10.1101/2025.07.31.664513

**Authors:** D. Campagner, J. Bhagat, G. Lopes, L. Calcaterra, A. G. Pouget, A. Almeida, T. T. Nguyen, C. H. Lo, T. Ryan, B. Cruz, F. J. Carvalho, Z. Li, A. Erskine, J. Rapela, O. Folsz, M. Marin, J. Ahn, S. Nierwetberg, S. C. Lenzi, J. D. S. Reggiani, SGEN group – SWC GCNU Experimental Neuroethology Group

## Abstract

Ethological behaviours are a powerful tool for neuroscience since they leverage the robust neural computations shaped by the species’ evolution to study the neural basis of cognitive functions. However, such behaviours are often transitory and dependent on factors that vary over space, time and number of individuals, making them difficult to capture with standard laboratory tasks. Here we present Aeon, an open-source platform designed for continuous, long-term study of self-guided behaviours in multiple mice and simultaneous recording of brain activity within large, customizable habitats. By integrating specialized modules for navigation, nesting and sleeping, escaping, foraging, and social interaction, Aeon enables the expression of key ethological behaviours while achieving experimental control and multi-dimensional quantifications from sub-millisecond to month-long durations. Its software architecture ensures robust data acquisition via many synchronized data streams and delivers a new standardised, unified data format that yields seamless, integrated analysis pipelines. Using assays such as digging-to-threshold and social foraging, Aeon reveals how mice adapt strategies in a changing environment and in response to conspecifics. Aeon bridges ecological relevance with rigorous experimental control to advance our understanding of how neural circuit activity gives rise to a range of highly conserved and adaptive behaviours.

## INTRODUCTION

Neural computations underlying behaviour have been sculpted over millions of years by the ecological niche species have evolved into^1,2^. By integrating ethological relevance into laboratory experiments, recent studies have exploited the tight link between behavioural ecology and neural circuit function^3–6^, leading to new insights into the biological implementation mechanisms of cognitive processes such as decision making, learning, and navigation.^3–9^

While these studies represent the first steps towards establishing a mechanistic neuroethological approach, reproducing ethologically realistic conditions under controlled laboratory settings remains challenging. Ethological behaviours are by their very nature unconstrained and multimodal^10^, taking place over large spatial extents, varying time scales, and dynamic social contexts (Fig. 1a). These factors critically shape behaviour and neural dynamics in most mammals^11–19^, including *Mus musculus*^20–23^, the predominant model organism in systems neuroscience.

**Figure 1.**
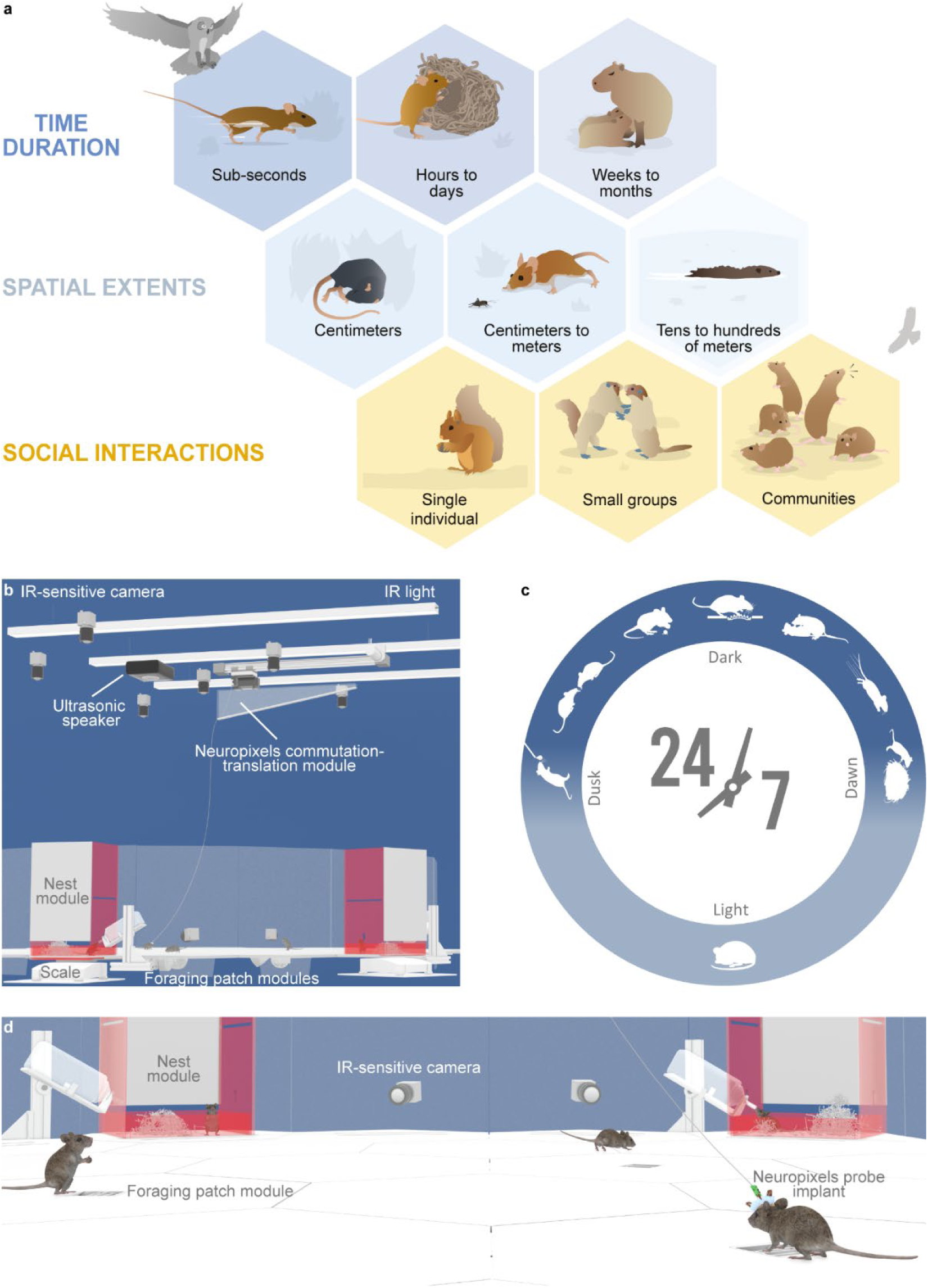
The Aeon project. **a.** Examples of natural behaviours in rodents that take place over varying timescales, spatial extents, and numbers of individuals (image credit: Jil Costa). **b** to **d**. Close-up views of an Aeon habitat from different vantage points (**b** and **d**) in which mice chronically implanted with Neuropixels probes and non-implanted mice can live while expressing a variety of natural behaviours including exploring, drinking, escaping, foraging, nesting, sleeping, eating and interacting socially (**c**). The figure depicts some of the core devices and modules that compose a habitat. Above the foraging floor: the overhead infrared LED lighting system, ultrasonic speakers, and the Neuropixels commutator-translation module. Embedded into or around the floor: the foraging patch modules, nest and scale modules, and surround IR cameras. Note: for visualization purposes we depict clear walls, but they are made of red-tinted see-through acrylic.

To encompass these complexities, an ideal experiment platform should meet several criteria in allowing animals to express a rich repertoire of natural and learned behaviours while supporting mechanistic neuroscience research. First, it should allow researchers to record and manipulate the behaviour of multiple animals simultaneously with high spatio-temporal resolution in relatively large and complex environments over extended periods of time. Second, it must permit control over experimental parameters, ensuring rigorous, reproducible experiments despite the inherently unconstrained nature of ethological behaviours. Third, it should support long-term physiology—particularly high-density neural recordings—while remaining modular and agnostic to behaviour type and assay structure, to accommodate the high dimensionality of ethological behaviours. Fourth, data acquisition, storage, interrogation and analysis must reflect the continuous nature of these experiments, requiring new standardized data formats and pipelines. Lastly, such a platform would ideally be modular and open source to enable opportunities for customisation, wide dissemination, collaboration, and reproducibility. Collectively, these approaches would allow the identification of generalisable principles of neural computation across specific neural circuits or brain regions that support a range of innate and adaptive behaviours.

To fulfil the requirements described above, we developed Aeon, a new platform to study the neural basis of ethological behaviours over time scales of weeks to months continuously (Fig. 1c) in multiple mice housed in large and modular habitats (Fig. 1b,d). Below, we detail the design and capabilities of the Aeon platform and showcase its use with results from long-term experiments during which we recorded the neural activity and behaviour of mice engaged in ethological assays, including social foraging.

## RESULTS

### Platform overview

Aeon provides an end-to-end experiment pipeline that streamlines habitat design, experiment control, data acquisition, data management, quality control, and analysis (Fig. 2). The platform is built upon two core principles: modularity and scalability. These principles are embedded throughout its design, from the physical assembly of habitats to the seamless integration of devices and the efficient handling of large-scale data, enabling researchers to tailor the platform to specific experimental needs.

**Figure 2.**
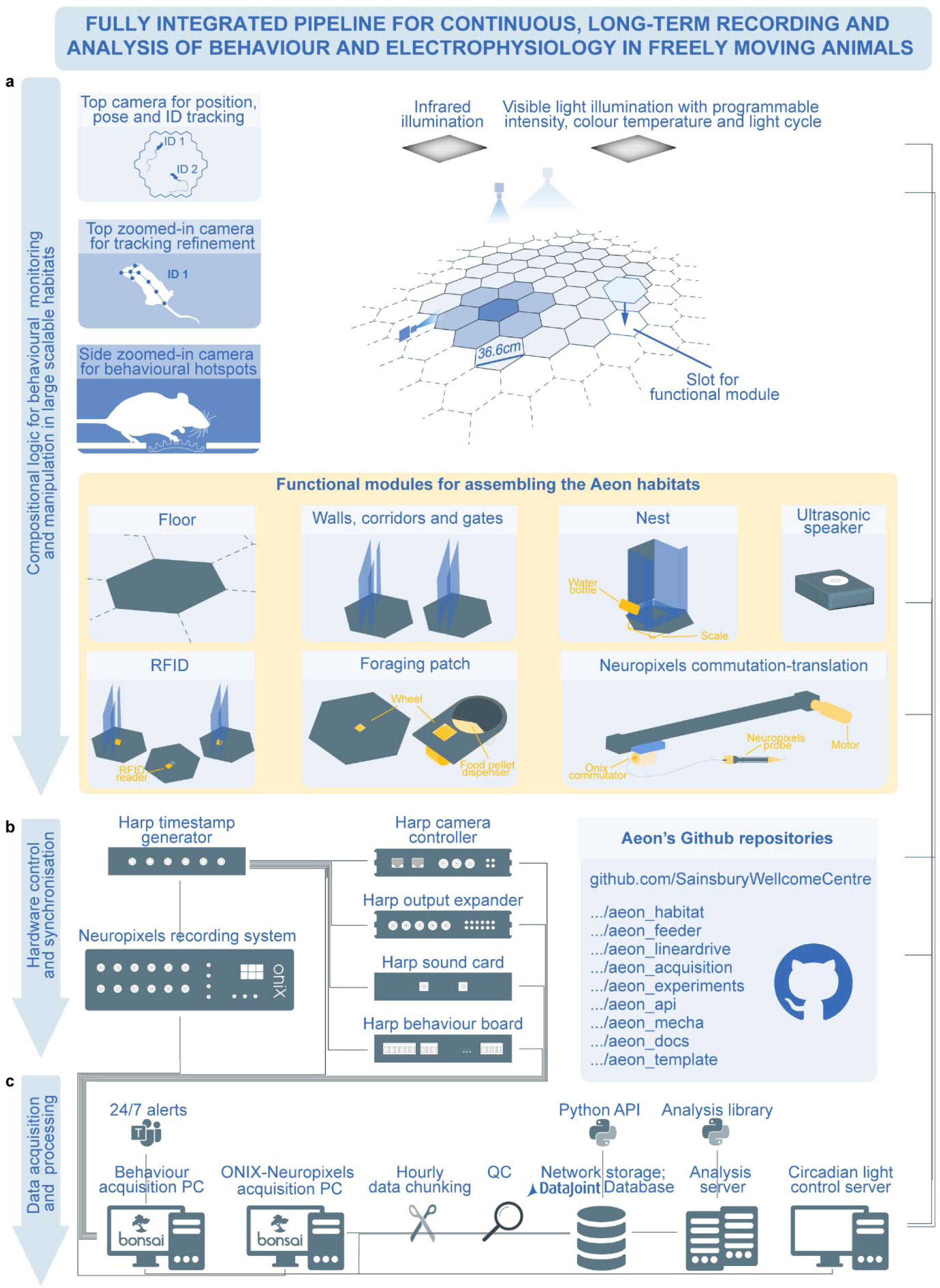
Aeon platform fully-integrated pipeline. **a**: Schematic illustrating the compositional hardware that forms an Aeon habitat. The Aeon platform incorporates a large variety of modules that can be assembled to form a honeycomb floor with a variety of functional configurations and extensions. Animal behaviour in an Aeon habitat is monitored by multiple infrared high-speed cameras with varying lenses serving three complementary purposes: 1) a full-field overhead camera for real-time subject identity, position and pose tracking; 2) zoomed-in overhead cameras for tracking refinement; 3) high-magnification cameras to monitor specific behavioural hotspots. The habitats are under constant infrared illumination and dynamic (both intensity and colour temperature) visible light illumination. **b. Left**: All Aeon devices are acquired and synchronised via Harp devices. The master clock for system-wide time alignment is provided by the Harp master clock generator. Neuropixels probe data are acquired with the OpenEphys ONIX acquisition system, which includes a PCIe card located in the ONIX-Neuropixels acquisition PC. **Right**: full list of the github repositories of the Aeon projects, containing hardware implementation (aeon_habitat, aeon_feeder, aeon_linear_drive), software implementation (aeon_acquisition, aeon_experiment, aeon_template), data processing (aeon_api, aeon_mecha) and documentation (aeon_docs, aeon_template). Links to the repositories and detailed explanations of their content can be found in the project website (https://aeon.swc.ucl.ac.uk/getting_started/repositories.html). **c.** Schematic illustrating data acquisition and processing. Data streams are acquired by a behavioural acquisition computer running Bonsai, which is responsible for controlling all aspects of an experiment. A second, neural acquisition computer, also running Bonsai, controls the ONIX system. Acquired data undergoes continuous quality checks, with alerts sent to researchers on a specified application interface (e.g. Microsoft Teams). Data is chunked at a fixed interval before being automatically moved to larger, long-term raw data storage. A set of Python routines is responsible for processing the data in this storage, and ingesting them into a DataJoint database. A final set of DataJoint Python routines are regularly and automatically called on the processed data to generate and perform standard visualizations and analyses during experiment uptime.

An Aeon habitat is built from modular, hexagonal tiles that accommodate a wide array of functional modules (12 modules/m^2^; Fig. 1b,d, 2a and Box 1). These modules define the spatial architecture of the habitat, provide nesting sites, water, or food, and support the study of ethologically relevant behaviours, such as escape and foraging (Fig. 1b and 2a). Many habitat configurations are possible via module rearrangement. Each module can be equipped with an RFID (Radio-Frequency Identification) reader for subject identification (Fig. 2b). Cameras with different degrees of zoom and frame rate are placed throughout the habitat to capture different aspects of behaviour while performing position tracking, subject identification and pose estimation (Fig. 2a). Additionally, many other devices, including weight scales and Neuropixels probes,^24,25^ have been integrated. Aeon recording devices use the Harp clock synchronization protocol^26^, which enables system-wide temporal alignment down to 50 us precision (Fig. 2b). In the Aeon platform documentation, we provide extensive instructions on how to assemble and control an Aeon habitat and each Aeon module (https://aeon.swc.ucl.ac.uk/user/aeon_modules.html#target-aeon-modules and https://aeon.swc.ucl.ac.uk/user/tutorials/general_experimental_workflow.html).

All behavioural data is recorded on a “Behaviour acquisition PC”, and all neural data on a separate “Neural acquisition PC” (Fig. 2c). Aeon employs an automated protocol in which data streams are written to files at a specified interval and transferred to long-term raw data storage (Fig. 2c). Automatic quality control and processing routines are applied to the data before ingestion into a MySQL DataJoint database^27^. Processed data used for most analyses can be queried from the database, while all raw data remains accessible via the low-level Aeon Python API, which is also called during database ingestion (Fig. 2c). To leverage the modular design of the habitat and data pipelines, experiment control must be precise, scalable, and adaptable. Aeon achieves this using Bonsai, an open-source, compiled, high-performance, reactive visual programming language designed for synchronous and asynchronous data acquisition and online processing^28^. To support the Aeon project, we developed several new Bonsai packages and workflows specifically for orchestrating dynamic habitat updates, interfacing with hardware, and logging all experiment data.

### Solutions to scientific and technological challenges in long-term neuroethology experiments

A defining strength of the Aeon platform is its ability to facilitate continuous behavioural experiments and neural recordings over extended periods—from weeks to months—while handling substantial data throughput. Achieving this goal required overcoming multiple challenges at the hardware, software, and experiment structure levels.

#### Time-alignment of many high throughput recording devices

In a typical experiment (for example the ones described in the section *Recording and quantification of social behaviours*), we record from over 30 unique hardware devices and over 60 unique data streams, and when including electrophysiology data from two Neuropixels probes, the total acquisition throughput can reach up to 200 GB per hour. It is therefore crucial to have a simple, robust method for online time alignment of all data streams at sub-millisecond precision, eliminating the need for extensive post-processing. We achieve this using the Harp clock synchronization protocol^26^: a single clock signal is multiplexed from a Harp clock synchronizer, via intermediary Harp devices such as output expanders, behavioural boards, and camera controllers, to all recording devices. This results in temporal alignment of all hardware data streams down to 50 us precision (Fig. 2b) and of all software data streams to milliseconds precision.

##### Box 1

**Aeon functional modules**

The Aeon platform supports a variety of functional modules designed to capture different aspects of ethological behaviour. Instructions on how to assemble these modules can be found in the project website: https://aeon.swc.ucl.ac.uk/user/aeon_modules/hardware.html#target-hardware. The list below includes only the modules used in the work presented here, but new modules will be made available at the link provided. See Fig. 2a for a pictorial representation of the modules. For technical specification see *Methods*.

###### Floor module

A plain, white acrylic hexagonal tile that forms open-field areas where mice can explore and interact.

###### Wall and corridor modules

These modules comprise a floor tile fitted with one or two parallel slots for red-tinted, transparent acrylic walls. By combining multiple units one can build barriers, partitions, long corridors or complex mazes.

###### The foraging patch module

The foraging patch module, consisting of the foraging wheel and the pellet delivery system, provides a means for animals to obtain food in the habitat by rotating the wheel, simulating a naturalistic digging action. The wheel’s pivot is supported by ball bearings for unidirectional rotation and a motion sensor (magnetic encoder) measures the wheel’s movement. Pellets are stored in an under-floor hopper and are dispensed onto the wheel via a motor-driven spinning disk as a function of the distance spun.

###### RFID module

The floor, foraging patch, walls and corridor modules can be equipped with an RFID reader that is mounted under the arena floor. This is used to read an RFID chip implanted in the mouse flank and enables animal identification.

###### Nest module

The nest is designed to provide a dedicated nesting site and access to water, while continuously monitoring the weight of the animals. A nesting box is attached to a scale for precise weight measurement and is surrounded by modified habitat outer walls. A water bottle can be suspended on the side of the nesting box, with its spout protruding into the nest through a wall opening. The nest is typically equipped with bedding materials and enrichment.

###### Ultrasonic speaker module

Allows playback of ultrasonic stimuli inside the habitat—for example, to elicit defensive behaviours.

###### Neuropixels commutation-translation module

To minimise the tethering cable’s interference with natural behaviour and prevent tangling during extended periods of recording, the Neuropixels commutation-translation system integrates the ONIX commutator with a servo motor-powered linear actuator, enabling closed-loop adjustment based on the animal’s 2D position and head direction.

#### Structuring, querying and analysing data across space, time and subjects

Another key challenge in continuous, long-term experiments is establishing an approach to yield structured, standardized data with minimal data loss. Storing data in large individual files is impractical for continuous, long-term recordings: a single hardware failure or file corruption could result in the loss of weeks or months of data. Additionally, a 65 TB dataset from a full two-week experiment (assuming a rate of 200 GB / hour) would be far too large for typical desktop storage drives, and individual files may not even fit on disk. Large files also make data querying challenging, as naively extracting specific time periods of interest requires loading the entire file into memory, which would be impossible given typical desktop memory capacity.

Furthermore, Aeon experiments do not align with conventional divisions into subjects, sessions and trials^29^. Different subjects may enter or exit the habitat multiple times within a single acquisition epoch, and the habitat facilitates spontaneous behavioural expression and self-paced interaction with environmental modules. Rules in the environment are designed for multi-subject interaction: initiation may be triggered or interrupted by any subject at any time. Existing approaches to standardising raw data schemas around normative principles such as subject, sessions or trials^29^ are thus not well suited. Instead, we adopted a structured storage specification organised around devices, time intervals (“chunks”), and continuous recording periods (“acquisition epochs”), both to efficiently manage high data volumes and allow robust long-term, flexible, and experiment-agnostic logging (Fig. 3a,b and Extended Data Fig. 1). In Aeon, all data streams belonging to an acquisition epoch are written out to individual files in chunks (by default, every hour), grouped by acquisition device. Within a chunk, data is organised via a globally referenced Harp timestamp, ensuring strict file time boundaries. This chunked, device-grouped file structure also facilitates periodic transfer of data from an acquisition PC to a larger network storage to avoid experiment interruptions due to insufficient local storage.

**Figure 3.**
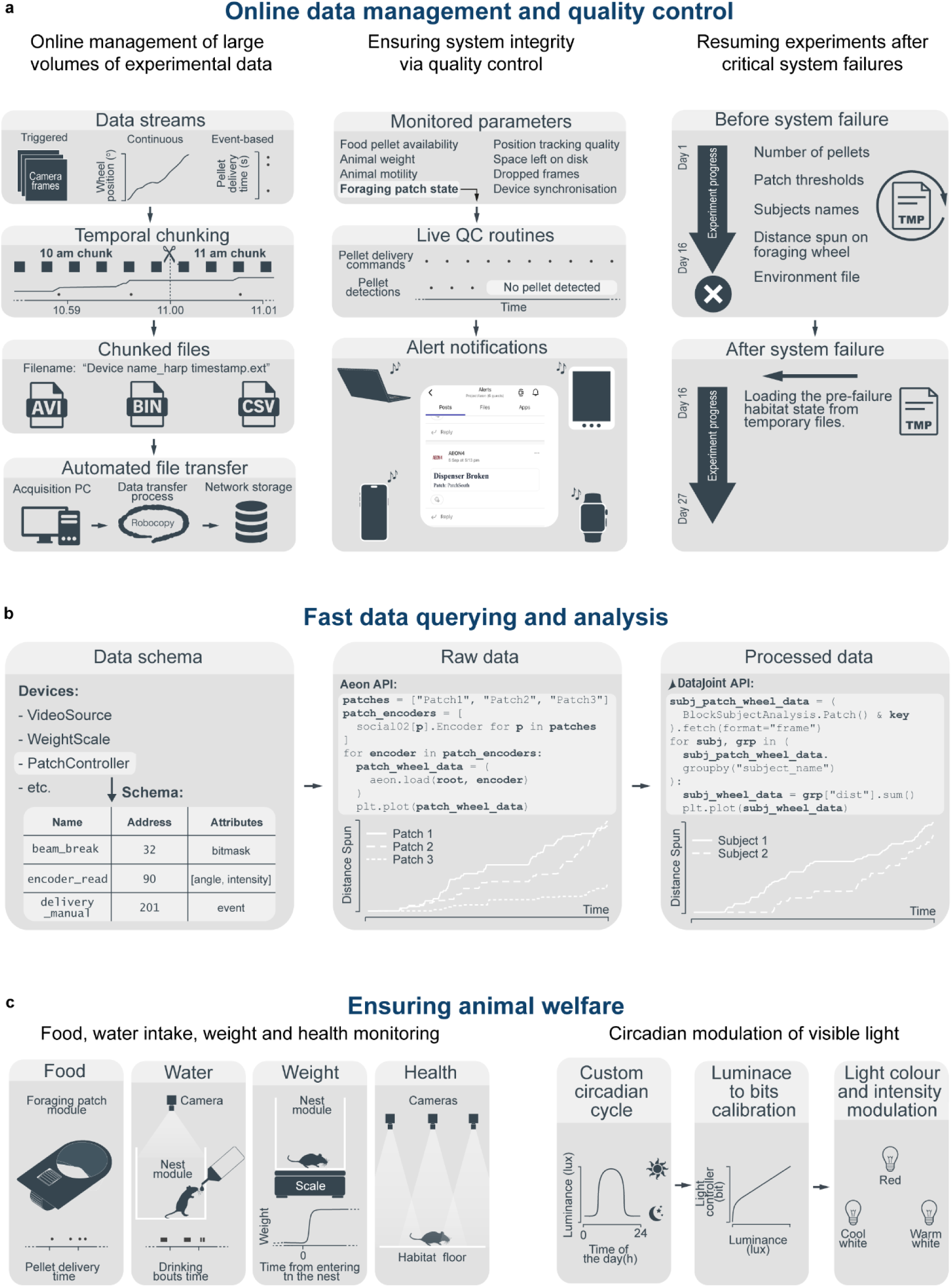
Scientific and technological strategies for long-term neuroethology experiments. **a. Left:** Aeon acquisition handles both continuously sampled and discrete, event-based data streams. At regular “chunk” intervals (by default every hour), data from each unique stream is written out to file, and all files in a chunk can be automatically transferred from the acquisition computer to a larger storage system. **Middle:** User-specified acquisition system information (including storage space and device synchronization status), animal information (including weight and motility), and data stream properties (including position tracking and foraging patch state) can be continuously monitored via live quality control (QC) routines. As a particular example, we highlight QC routines for foraging patch module state, in which an alert is sent when no pellet detection event follows a pellet delivery command, indicative of a foraging patch module failure. **Right:** Recovery from critical system failures (e.g. power or operating system failures) is handled via temporary files. During experiment runtime, Bonsai saves relevant experiment logic parameters to temporary files. After a system failure and subsequent experiment workflow restart, the temporary files can be optionally read from to restore the experiment state to its immediate pre-failure status. **b.** Aeon data architecture sets standardized data schemas to enable fast data querying and analysis. All data streams are grouped into specific devices, and each data stream within a device is given a unique name and register address (**left).** The low-level Aeon API provides Python classes that correspond to raw data streams. Class objects are passed to the load function, which retrieves the data stream’s name and register address to locate and read the corresponding files. As a particular example, we highlight encoder data streams in patch devices: patch names, which must match file directory device names, are specified in patches, a list of Encoder objects corresponding to each encoder’s respective patch is created from patches, and the raw data corresponding to each encoder can be read via ‘load’ and plotted (**middle**). Via the high-level DataJoint API, processed data can equally be easily queried and plotted. In this example, subject-specific processed data, stored in the BlockSubjectAnalysis.Patch table, is filtered by a key that specifies a particular time window. As shown, in just a few lines of code, the data from this table can be grouped by subject, aggregated across patches, and plotted, to visualize the total distance foraged (summed over all patches) per subject. (**right**). **c.** Many parameters relevant to animal welfare are monitored continuously throughout Aeon experiments. In particular, Aeon tracks each animal’s food consumption and weight, along with available food levels and foraging patch module status. Water consumption and animal health status can be monitored via the array of cameras that capture a habitat **(left)**. Additionally, a customizable light cycle can be programmatically defined and continuously monitored to ensure that habitat lighting conditions remain consistent with experiment expectations **(right).**

Adopting this directory structure allowed us to design a general API for directly querying data across experiments and time ranges. (Fig. 3b and Extended Data Fig. 2). DataJoint database routines can leverage the low-level API to process raw data into analysis-ready tables and automatically ingest them into a database on a periodic basis. This enables researchers to explore preliminary results in real-time while an experiment is still in progress (Fig. 3b and Extended Data Fig. 3). In the Aeon platform documentation, we provide Python tutorials for accessing data directly using either the low-level API or DataJoint (https://aeon.swc.ucl.ac.uk/user/how_to.html#target-how-to).

#### Continuous, long-term quality control and experiment continuity

In long-term experiments, maintaining consistent and reliable data over days to weeks is critical, as small errors from hardware failures and/or data corruption can accumulate and ultimately jeopardize entire datasets. To mitigate this risk, the Aeon platform incorporates a comprehensive suite of automated quality control checks. These include detecting dropped camera frames, failures in pellet deliveries, and unexpected deviations in time alignment across all recording devices. To ensure animal welfare, the system also tracks individual subject weights, pellet dispenser levels, and overall behavioural activity. Lastly, it safeguards data acquisition by monitoring disk storage and memory usage, network drive connectivity, and many other potential critical failure modes which might compromise experiment data and control (Fig. 2c and Fig. 3a).

In the first instance, quality control and processing routines are applied online to individual data streams. Any detected abnormality is immediately sent via a Bonsai webhook to a customizable app (in our case Microsoft Teams). Additionally, any experiment state defined as persistent is regularly saved to temporary files, allowing the experiment protocol to be resumed in the event of a critical system failure (Fig. 3a).

In the second instance, each raw data stream chunk file can be processed for data corruption and inconsistencies before ingestion into a DataJoint database. While we cannot correct fundamental data errors at this stage, it is often possible to interpolate missing data, reprocess the output of specific algorithms such as pose tracking, label specific time ranges as invalid or unobserved, and perform other important data-cleaning operations (see *Methods*). Importantly, the raw data remains unaltered; all processing yields separate ‘processed’ data files.

#### Ensuring Animal Welfare During Extended Experiments

The 24/7 nature of Aeon experiments requires systems that safeguard animal welfare (Fig. 3c). These systems ensure uninterrupted access to food, water, and nesting sites, while also enabling continuous monitoring of each subject’s weight.

Food is provided through the foraging patch modules, which dispense 20 mg pellets (Fig. 1b,c, 2 and Box 1). Each pellet delivery is logged to quantify the food intake for each animal. Nesting sites in an Aeon habitat are provided by the nest modules (Box 1). A high-speed camera focused on a bottle spout fit into the nest’s wall allows monitoring of water consumption (Fig. 2b and 3c). The nest module’s floor serves as the load plate of a scale, enabling weight measurements each time a mouse remains briefly motionless in the nest (Fig. 3c and Extended Data Video 1). Lastly, the overall health status of each mouse can be monitored in real time via the multiple high-speed cameras that provide full habitat coverage (Fig. 2 and 3c).

The long-term housing of mice in Aeon habitats also requires precise control of environment luminance throughout the day to replicate diurnal cycles. The Aeon platform supports the programming of luminance patterns that change throughout the day (Fig. 2 and 3c), with single-minute temporal resolution. To modulate spectral composition as a function of the time of day or specified environmental conditions, the lighting system combines cool white (4000 K) and warm white (3000 K) fluorescence light sources, and red LEDS.

#### Enforcing standards

As experiments may run continuously for months, the lack of a well-defined system for enforcing metadata standardization, experiment documentation, and version control can lead to issues with data integrity, reproducibility, and cross-experiment comparability: over time, datasets may become difficult to interpret, integrate, or analyse. Moreover, as hardware configurations and experiment logic evolve, it is essential to have mechanisms to dynamically update databases, maintain structured documentation, and ensure software reliability without introducing inconsistencies or eschewing backwards compatibility. To address these challenges, we introduced data contracts, which link experiment control, data acquisition, data storage, and data analysis to enforce standardization throughout the Aeon pipeline.

At experiment onset, a Python module is called to parse a Bonsai workflow XML file and automatically generate a YAML metadata file which captures a snapshot of all the experiment’s relevant habitat parameters and device metadata. This metadata file lives within the raw data directory that corresponds to the experiment. After the first experiment chunk, separate Python code can then parse this metadata file to populate existing Datajoint database tables.

Additionally, when required, new database tables can be automatically generated using Python source code generators, ensuring that devices listed in the metadata file without existing database tables are seamlessly integrated.

Documentation for the entire Aeon project is managed through the aeon-docs GitHub Pages repository, which incorporates other Aeon repositories (Fig. 2b) as submodules, enabling centralized access to all Aeon resources and automated generation of new documentation whenever any of the modules in these repositories is updated.

### Ethological behaviours in Aeon habitats

A key requirement of the platform is that subjects living in an Aeon habitat retain essential features of their behavioural repertoire. To test this, we placed mice in an Aeon habitat equipped with three foraging patches, one ultrasonic speaker and one nesting sites for multiple days. We observed that mice spontaneously exhibit sleeping, drinking, eating, nesting, escaping, social interactions and other ethological behaviours typical of the species (Fig. 4, 6b and Extended Data Video 2 - 8; total of 19 mice).

**Figure 4.**
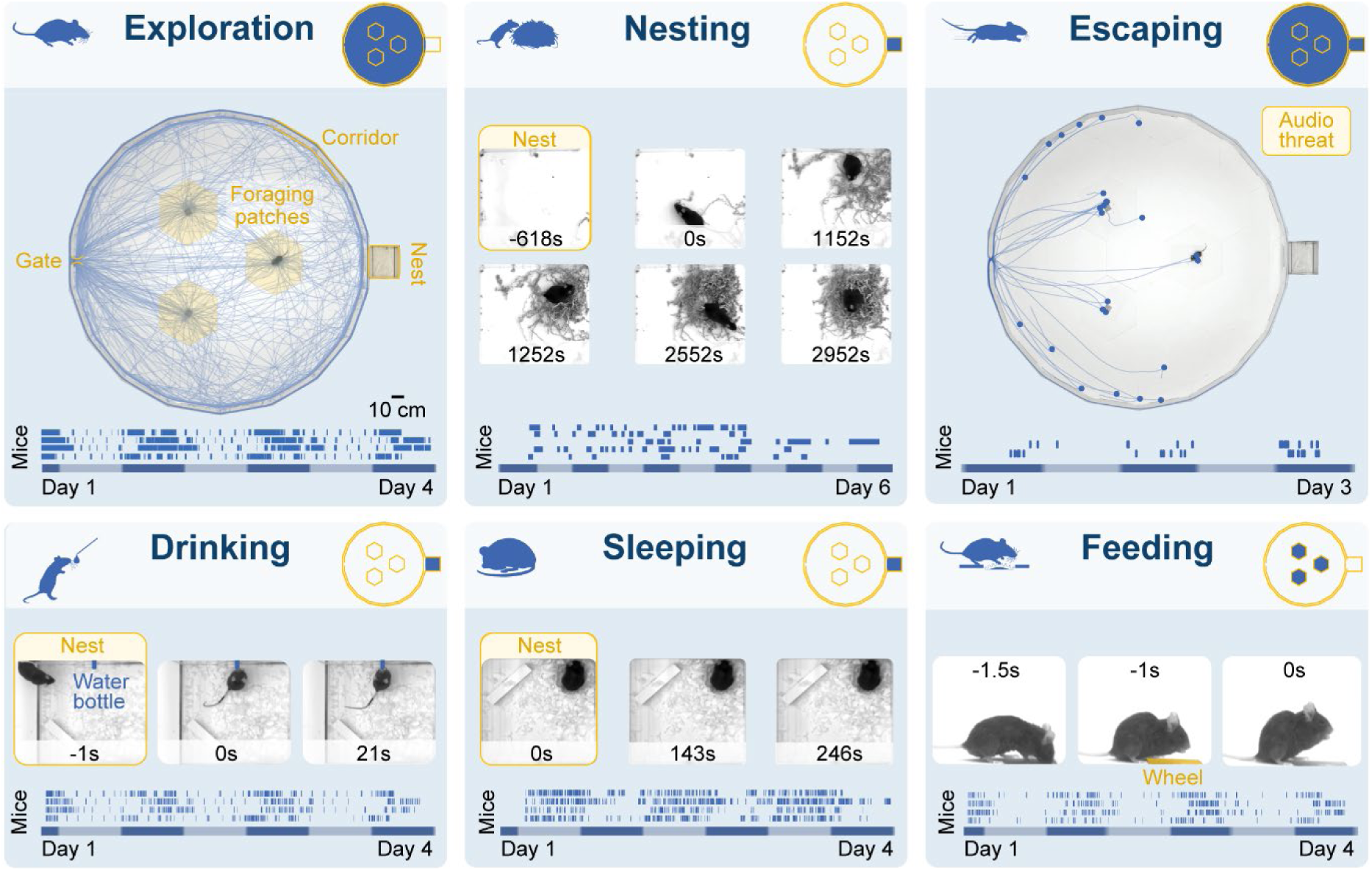
Ethological behaviour in an Aeon habitat. Examples of natural behaviours extracted from continuous, multi-day video. Plotted blue ticks represent the behavioural events. On the x-axis, light-dark luminance cycle: blue bars correspond to low luminance periods, while grey bars correspond to high luminance periods. Insets in the top right corner of each panel are schematics of an Aeon habitat: here blue areas indicate the regions where the respective behaviour was measured. (see Exploration panel for details on habitat components).

When housed in a habitat with 11 hours of high luminance (75 lux, light condition), 11 hours of low luminance (1 lux, dark condition) and two, 1 hour blocks of intermediate luminance (40 lux, dawn/dusk)^30^, mice synchronised their behavioural patterns to the habitat’s cycle in a way characteristic of nocturnal animals: exploration bouts were predominantly observed during the low-luminance period (exploration during dark period: 90.1%; exploration during light period: 9.9%), while sleeping episodes occurred primarily during the high-luminance period (sleep during dark period: 22.3%, sleep during light period: 77.7%). During exploration, mice engaged in foraging behaviours by operating the foraging patch modules available in the habitat (Box 1; Extended Data Video 6 - 8). Mice spontaneously recognise the nest module as their nesting site, gathering nesting material and building nests within (Fig. 4 and Extended Data Video 2).

Mice not only engaged in spontaneous behaviours but also expressed adaptive behaviours including escaping to safety from auditory threats (reaction time: 443 ±207; peak speed: 1.4±0.09 m/s; escape probability: 0.75; Fig. 4 and Extended Data Video 4) and various social interactions (Fig. 6 and Extended Data Videos 10 - 12).

Taken together, these findings demonstrate that an Aeon habitat provides an ecologically valid environment in which mice can naturally express a broad repertoire of species-specific behaviours.

### Parametric ethologically relevant assays in Aeon

The Aeon platform not only provides tools to monitor and measure mouse ethological behaviours but is also equipped with modules to parametrically manipulate them and test hypotheses experimentally.

One such behaviour is foraging, a powerful framework for decision making in natural settings^31,32^, which can be studied in Aeon habitats via the foraging patch module (Fig. 5a and Box 1). When interacting with this module, which serves as the sole source of food in an Aeon habitat, mice must continuously gauge effort and reward while performing a parametric digging-to-threshold foraging assay, in which they simulate digging by spinning an underground wheel using their forepaws (Fig. 5a, b and Extended Data Video 6 and 7). A 20 mg food pellet is automatically delivered on the wheel by an underground pellet dispenser when a certain spinning distance (pellet threshold) is reached. The rules governing threshold changes can be arbitrarily specified (Extended Data Fig. 4). During this assay, mice must balance energy expended from wheel spinning with the amount of reward obtained, through consumption of one or more food pellets.

**Figure 5.**
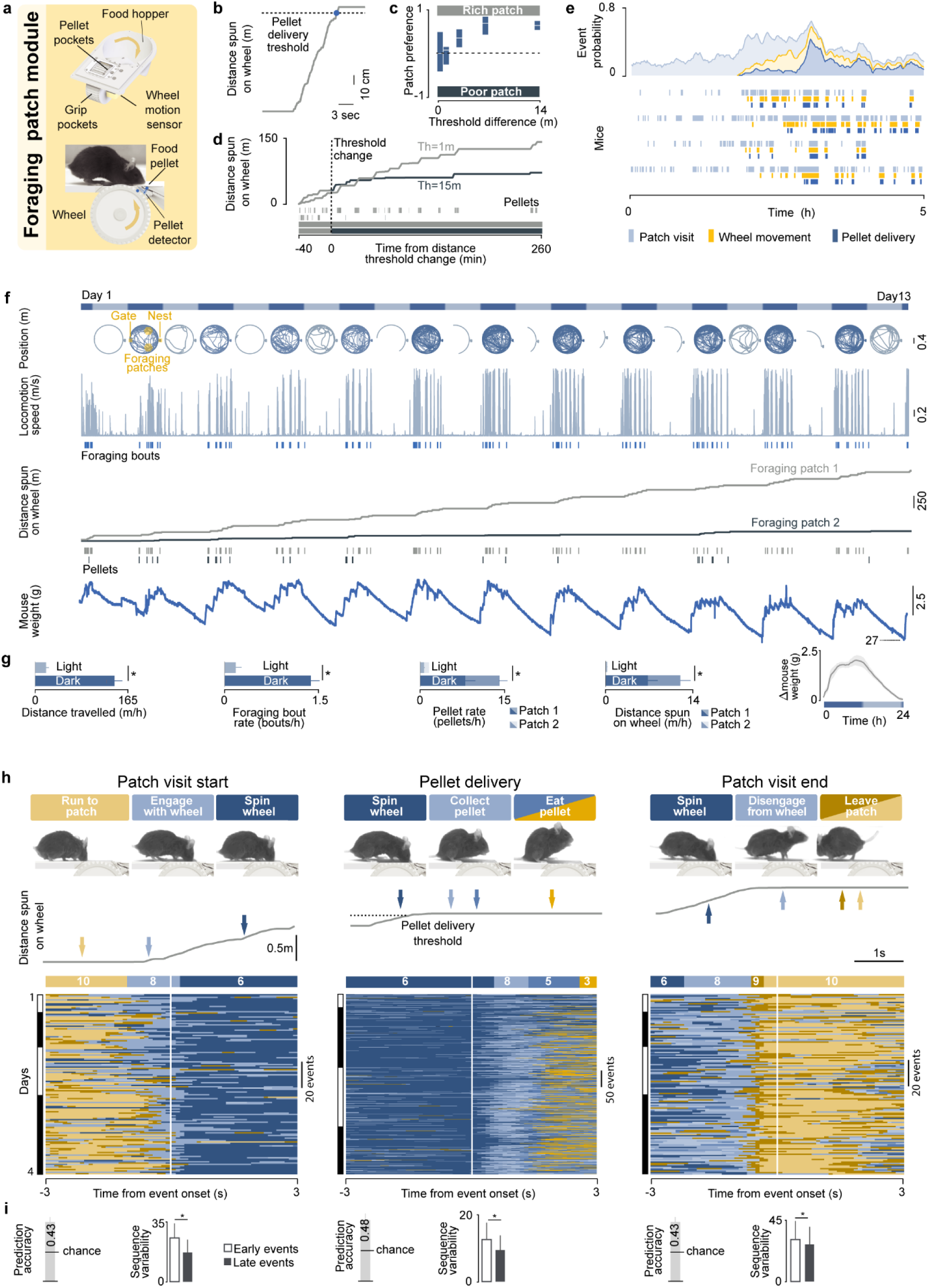
Characterisation of foraging behaviour over multiple timescales. **a.** 3D model of the foraging patch module **b.** Example of one foraging episode during which the mouse had to spin the foraging module wheel to a threshold of 1 m to get a reward of one pellet. **c.** Patch preference computed from experiments like the one illustrated in **d.** Each experiment was performed in an arena with two foraging patches. At the start of the experiment, both patches present a distance threshold of 1m. After 20 pellet deliveries jointly from both patches, the patch with the highest pellet count switched threshold to a new value that could range between 1 m and 15 m (poor patch, 15 m in d), while the other stayed the same (rich patch). Box plot: median, IQRs. N= 6 mice. **e.** Data from 4 mice showing the first time each interacted with the foraging patch. **Bottom**: raster plot of the following events grouped by mouse, patch proximity, wheel movement, or pellet delivery. **Top**: averaged event probability (bin size = 15 min) **f.** Example of a two-week-long, two-patch, single-mouse foraging assay (patch threshold 1m in both patches). The habitat configuration is similar to the one depicted in Fig. 4, but with two foraging patches instead of three. The bar at the top indicates periods of high (gray) and low (blue) environmental luminance. Traces below indicate the x,y position of the mouse during the respective period (light or dark). Golden shaded areas show the location in the habitat of the foraging patches, nest box, and gate to access the habitat centre. **g.** Global statistics from 5 mice performing the two-patch foraging assay over multiple days (* p<0.05). **h.** Top: Examples from the 10 HMM states describing different interpretable syllables in the foraging sequence, aligned to salient events in the foraging sequence, such as start of a patch visit, pellet delivery, and end of a patch visit. Dark blue to brown shades indicate different HMM states. Arrows indicate the time point in which the frame was acquired, and are coloured according to the classified HMM state given the frame. Bottom: state sequence aligned to the respective foraging event during a 4-day long foraging assay. Top: dominant state sequence and respective state number (see methods). **i.** Quantification of accuracy of predicting the occurrence of foraging events, based on behavioural sequence composition and sequence variability between early and late events across time (*, t-test, p-value<0.05, 4 mice) Error bars: standard error of the mean.

Mice can learn to operate the foraging patch module spontaneously, within hours from first exposure (146 ± 14 min; N=4; Fig. 5e). Differences in pellet threshold between patches parametrically controls mouse behaviour: when presented with two foraging patches that differ in pellet threshold, mice develop a preference for the patch with the lowest threshold (rich patch), and the strength of this preference scales with the magnitude of the difference between the two patches (Fig. 5c,d). Additionally, when multiple foraging patches are available and one is unrewarded(Extended Data Fig. 7a), mice learn to mostly avoid the unrewarded patch and forage from the others (Extended Data Fig. 7b). Mice consume on average 15 pellets per hour while performing this assay, resulting in hundreds of foraging episodes per day (Fig. 5g).

### Quantification of ethological behaviours over multiple time scales

The Aeon platform captures snapshots of the global descriptive statistics of habitat exploration patterns, interactions with foraging patches, and other behavioural variables over ethologically relevant timescales of weeks at millisecond resolution (Fig. 5f; Extended Data Video 8).

As noted above, for spontaneous ethological behaviours, foraging behaviour displays slow periodic oscillations phase-locked to the diurnal cycle of the habitat, with more foraging bouts performed and more pellets consumed during the dark phase. Such circadian oscillations are also reflected in biometric measures such as locomotion, sleep, and body weight (Fig. 5f,g; Extended Data Video 8).

A unique feature of the Aeon platform is that it allows pairing the above holistic descriptions to detailed quantifications of the fine structure of mouse behaviour over the entire experiment duration. To achieve this, we fitted Hidden Markov Models (HMMs) to animal kinematics and pose tracking data, and isolated short model states (state duration: median ± IQR 0.6 ±0.9) representing interpretable behavioural syllables. For example, we observed states such as engaging with the foraging wheel, spinning the wheel, and collecting a food pellet (Fig. 5h and Extended Data Video 9). These syllables were organised in behavioural sequences predictive of foraging events, such as the start of a foraging bout, the conclusion of a foraging bout, or a pellet delivery (Fig. 5h,i). The reliability of syllable transitions and sequences allows the prediction of foraging events over several days (Fig. 5i), and the sensitivity of our model is such that it can also capture subtle changes in sequence composition. We observed that behavioural sequences were less reliable in initial foraging trials, and that sequence reliability increases and stabilises over days (Fig. 5i). This may suggest that mice tend to attempt multiple behavioural strategies while learning to interact with a foraging patch module before converging to a more stable action policy set.

Together, these results demonstrate that the Aeon platform is capable of monitoring and manipulating foraging behaviour for several weeks to months, capturing behavioural motifs that span timescales from hundreds of milliseconds to weeks.

### Recording and quantification of social behaviours

Social context greatly influences rodent behaviour and neural computation^18,19,21,22,33^. To investigate this, we equipped the Aeon platform with technology that enables multi-animal experiments by tracking subject identity with high accuracy. Aeon achieves this via two complementary methods: event-based subject identification at behavioural hotspots using RFIDs, and continuous computer vision identity recognition using the SLEAP software package^34^.

#### RFID system

The RFID approach involves implanting each mouse with a miniature RFID microchip that emits a unique identification tag. RFID readers, positioned strategically within the habitat, reliably detect these microchips (detection probability: 0.83 ± 0.03) when the mouse is in close proximity (3.6 ± 0.2cm; Fig. 6a), within a few hundred milliseconds (detection latency: 0.23 ± 0.01 s). RFID readers were placed under the floor at specific locations: including at entrances to the nest, foraging patches, and entrances from the corridor to the habitat centre (gates). (Fig. 6a and Extended Data Fig. 7a).

**Figure 6.**
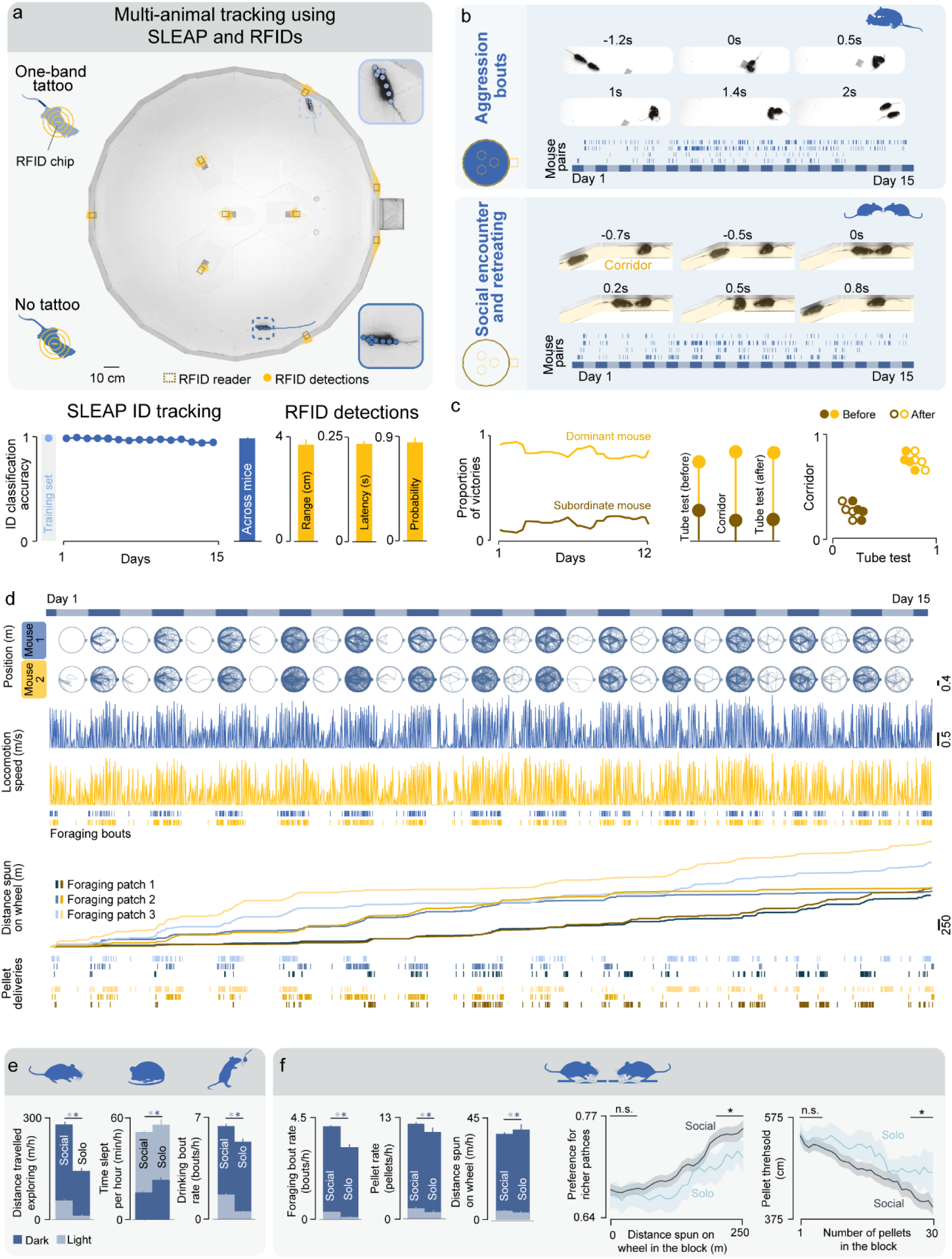
Multi-animal experiments in the Aeon ecosystem. **a. Top:** Overhead camera frame from an Aeon habitat with the following modules: four foraging patches, three gates, and one nest. Each of these modules was equipped with RFID readers (brown dotted rectangles) for animal identification. The location at which the RFID chip was detected by the reader is indicated as a golden dot. Shades of blue indicate the identity of the mouse assigned by SLEAP. SLEAP estimates of body part locations are overlaid onto each mouse. Left side insets: pictorial representation of mouse tail tattoos and implanted RFID chips. Right side insets: zoomed in image of each mouse. **Bottom left:** prediction accuracy of a SLEAP model in the training set (light blue) and testing set over the course of 15 days of experiment for one particular mouse, and for a population of 12 mice. **Bottom right:** quantification of RFID accuracy across mice (n=12). **b.** Examples of natural social behaviours extracted from continuous, multi-day video. Plotted blue ticks represent the behavioural events. On the x-axis, light-dark luminance cycle: blue bars correspond to low luminance periods, while grey bars correspond to high luminance periods. Insets in the top right corner of each panel are schematics of an Aeon habitat: here blue areas indicate the regions where the respective behaviour was measured. (see Extended Data Fig. 8a for details on habitat components). **c. Left**: continuous quantification of mouse dominance based on the proportion of victories during social encounter and retreating events over the course of a 12-day experiment. **Middle:** proportion of victories for the mouse in the lefthand panel during the social encounter and retreating events and during standard tube test performed before and after the experiment. **Right:** comparison between the proportion of victories during the standard tube test and social encounter and retreating events across four pairs of mice. **d.** Example of a 15-day, three-patch foraging assay with two mice. The habitat configuration is described in Extended Data Fig. 8a. The bar at the top indicates periods of high (gray) and low (blue) environmental luminance. Traces below indicate the x,y position of the mice during the respective period (light or dark). Locomotion speed, timing of foraging bouts and pellet deliveries, and time series of distance spun in each foraging patch are indicated in shades of blue for mouse one and shades of gold for mouse two. **e.** Differences between solo and social conditions across different ethological behaviours including exploration, sleeping and drinking in light and dark conditions across 10 mice. * p<0.05, t-test. **f. Left:** Differences between solo and social conditions in multiple foraging behavioural metrics (* p<0.05, t-test,) across 10 mice. **Right:** Metrics illustrating faster learning progression within each block during social conditions, as compared to solo conditions, in the same subjects (slopes of linear regressions for preference for richer patches versus distance spun on wheel in the block: solo = 0.065 ± 0.007, social = 0.094 ± 0.006; slopes of linear regressions for pellet threshold versus number of pellets in the block: solo = −2.98 ± 0.21, social = −4.46 ± 0.10; for all slopes p < 0.05). *: p < 0.05, t-test in the range indicated by the black line. Shaded areas and error bars: standard error of the mean. n.s.: not significant.

#### Computer vision tracking

For continuous, online, multi-animal tracking of pose (Extended Data Fig. 5) and identity, we used the computer vision software SLEAP, which performs inference using Convolutional Neural Networks trained on labelled data (Fig. 6 and Extended Data Video 10). For maximum accuracy, SLEAP identity models were trained both on zoomed-out and zoomed-in top view cameras frames. During inference these positions were mapped back to the full-view coordinate space of the habitat (Extended Data Fig. 6). To enhance individual identifiability, distinct band patterns were tattooed on mouse tails (Fig. 6a and Extended Data Fig. 6), providing a clear and stable marker that a trained SLEAP model can reliably use over weeks (Fig. 6a; mean classification accuracy: 0.96 ± 0.01).

#### Quantification of spontaneous social behaviours and social hierarchy

The resulting data enables automated or semi-automated classification of a large variety of social behaviours, including aggression bouts and social encounters and retreats (Fig. 6b, Extended Data Video 11 and 12). The latter can be used to infer hierarchical relationships over an experiment, generating continuous temporal rankings from most subordinate to most dominant individuals (proportion of victories is larger in dominant versus subordinate mice, binomial test, p<0.03; Fig 6c). These automated rankings are cross-validated against standard tube test assays^35^ (proportion of victories is larger in dominant versus subordinate mice, binomial test, p<0.02; Fig 6c), confirming their robustness (proportion of victories during standard tube test assay equal the one obtained with the automated ranking in all mice, Wilcoxon signed-rank p=0.25; Fig 6c).

#### Social context shapes ethological behaviours and learning

We next leveraged Aeon to test whether ethological behaviours are affected by social context. We conducted experiments that lasted approximately four weeks, in which pairs of mice were housed in an Aeon habitat and alternated living in periods of isolation and cohabitation. Throughout the experiment, mice foraged from three patches whose mean pellet thresholds - and thus relative resource availability - were dynamically updated every one to three hours (“blocks”) (Extended Data Fig. 7a-f provides full details of the assay). To maximise food intake, the mice needed to determine the “rich” patch (the patch with the lowest mean pellet threshold) during each block of time. We found that several ethological behaviours differed between solo and social conditions, across dark and light cycles. When housed together, individual mice showed greater exploratory drive, covering longer distances (distance travelled during dark: solo = 135.10 ± 3.63 m/h; social = 283.67 ± 3.74 m/h; p-value = 2.04e-151, t-test), and displayed more varied sleep (time slept during light: solo = 49.52 ± 2.86 min/h; social = 45.40 ± 1.59 min/h; p-value = 1.77e-202, t-test; sleeping bout duration during light: solo = 47.81 ± 2.60 min; social = 11.38 ± 0.79 min; p-value = 2.11e-45, t-test) and drinking patterns (drinking bouts rate during dark: solo = 5.42 ± 0.20 bout/h; social = 6.46 ± 0.16 bout/h; p-value = 6.68e-5, t-test; Fig. 6e and Extended Data Fig. 7). Foraging behaviour also differed: mice engaged in more foraging bouts in the social condition (foraging bouts rate during dark: solo = 3.27 ± 0.19 bouts/h; social = 4.22 ± 0.09 bouts/h; p-value = 7.74e-6, t-test), but each bout was shorter in duration (foraging bout duration during dark: solo = 3.40 ± 0.06 min; social = 2.93 ± 0.03 min; p-value = 3.23e-12, t-test; Extended Data Fig. 7 g), and they obtained more pellets overall together than when housed alone (pellet rate during dark: solo = 11.41 ± 0.59 pellet/h; social =12.50 ± 0.23 pellet/h; p-value = 0.009, t-test). Despite the higher activity and pellet yield, the total distance spun on the foraging wheel during social periods was shorter than during solitary periods, indicating more effective foraging in the presence of a conspecific (distance spun on wheel during dark: solo = 41.08 ± 2.46 m/h; social = 39.06 ± 0.81 m/h; p-value = 0.044, t-test; Fig. 6f left). Consistent with this, mice located the richer patches faster within each block during social foraging (Fig. 6f right), demonstrating a pronounced interaction between social context and learning under conditions of uncertainty.

### Long-term neural recordings

The Aeon platform not only enables the longitudinal characterisation of mouse behaviours over multiple time scales in large environments, but also permits concomitant monitoring of the underlying brain activity.

A key technique available in the platform is large-scale electrophysiological recordings using Neuropixels probes. Recording continuously for weeks in multi-meter enclosures required overcoming several technical challenges. The tethering cable had to minimize interference with natural behaviour and avoid tangling over the course of days and weeks. Additionally, the cable needed to accommodate exploration of a large habitat without creating excessive slack when the mouse was near the centre (Fig. 7b). We developed a commutation-translation system by integrating the ONIX commutator^36^ with a servo motor-powered linear actuator, enabling closed-loop adjustment based on the mouse’s 2D position and heading direction (Fig. 7a,b, Extended Data Fig. 8 and Extended Data Video 13). The combination of these two approaches enabled prolonged recordings (Extended Data Video 14) with virtually no changes in movement kinematics or exploration patterns across days (Fig. 7c, 8b and Extended Data Video 13). Following implantation of Neuropixels probes, natural behaviours, including foraging, remained unchanged and were synchronized to the habitat’s diurnal cycle. Implanted mice exhibited comparable numbers of pellets foraged, distance spun on the wheel, and distances travelled per hour with non-implanted mice (Fig. 5g and 8a,b). As expected, slow-wave sleep predominantly occurred during the habitat’s light phase, and more sporadically during the dark phase, with dynamics strongly anticorrelated with mouse movement (Pearson’s correlation coefficient −0.7, Fig. 8a,b).

**Figure 7.**
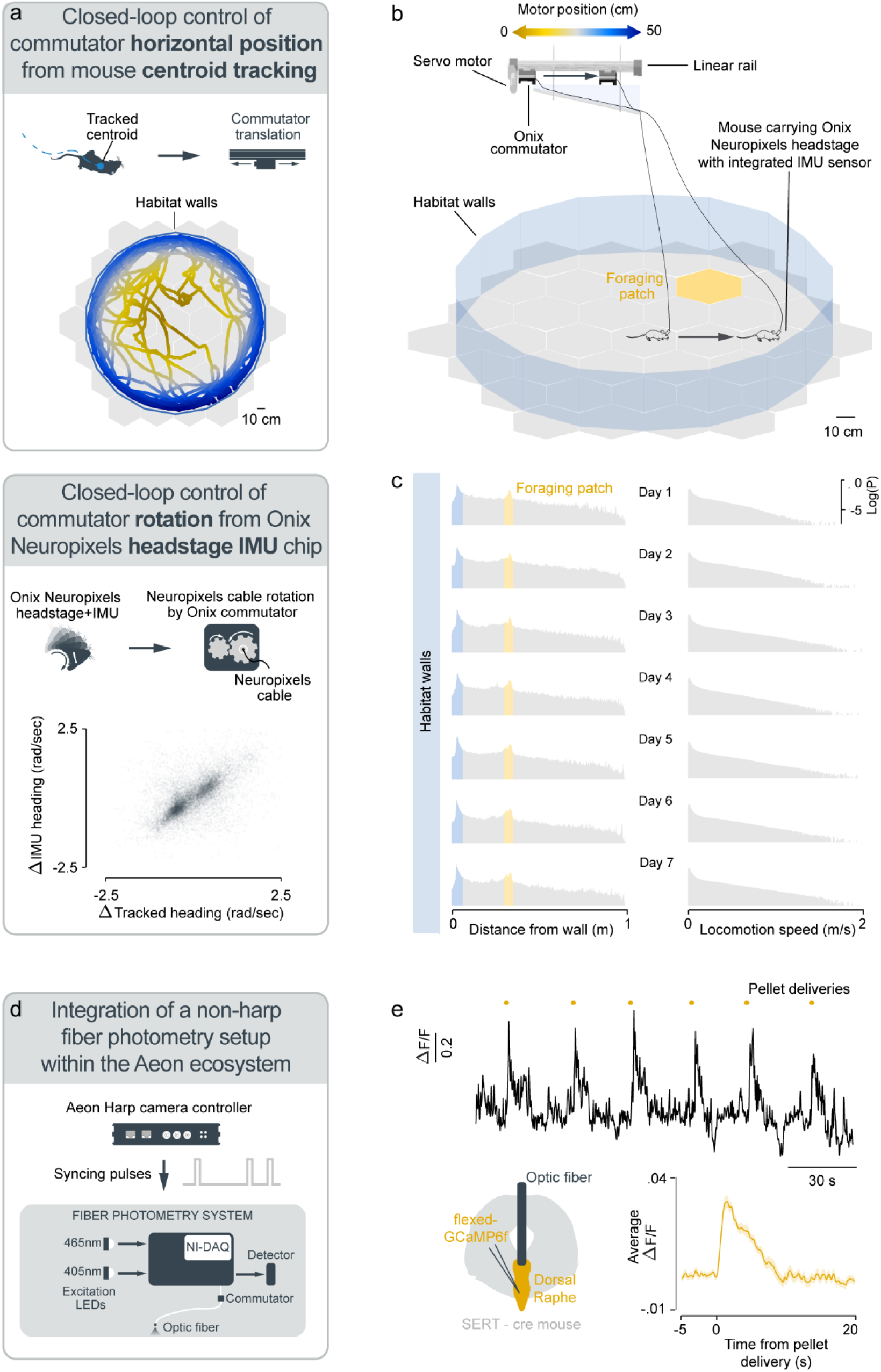
Approaches to record neural activity in Aeon habitats. **a. Top:** Schematic representing the approach to adjust the tether cable length as a function of the mouse location in the habitat: the ONIX commutator is mounted on a linear rail driven by a servo motor (Fig. 7a,b). This setup dynamically adjusts the tether length based on the mouse’s position within the habitat. A Bonsai workflow continuously tracks the mouse’s location in the habitat such that when the mouse moves toward the centre of the habitat, the rotary joint retracts along the rail to shorten the cable; conversely, when the mouse approaches the habitat’s edges, the rotary joint advances to release the cable. The mouse trajectory is coloured as a function of the linear motor position (see colour bar in panel **b)**. **b.** Bottom: Schematic representing tether tangle management using the ONIX commutator system. The ONIX system allows large-scale electrophysiological recordings using Neuropixels probes and features a custom-designed Neuropixels head-stage equipped with an integrated inertial measurement unit (IMU). The IMU tracks changes in the mouse’s heading direction, and this data is used to control the ONIX commutator, which untangles the Neuropixels tether in real-time based on mouse rotations. By utilizing the IMU output to guide the commutator, the system eliminates the need for a mechanical coupling between the commutator and the mouse’s head, resulting in a lightweight and highly flexible tether that minimizes stress on the animal. The plot shows heading estimates from the IMU sensor mounted in the Neuropixels head-stage versus the estimate derived from simultaneous videography of the tethered mouse exploring the habitat. **c.** Pictorial representation of a habitat during a tethered recording with Neuropixels probes. Arrows indicate the direction of movement of the mouse and the commutator. When the mouse is in the centre of the habitat, the linear motor moves the commutator backwards and the tether is shortened, and when the mouse moves away from the habitat centre, the linear motor moves the commutator forward, elongating the tether. **d.** Per-day distributions of distance from the closest wall and locomotion speed during a week-long recording session in the habitat illustrated in panel **b**. **e.** Schematic depicting the approach to perform fibre photometry recordings in Aeon habitats by coupling a non-harp compatible fibre photometry recording system to Aeon. **f.** Calcium bulk fluorescent signal recorded from a population of serotonergic neurons in the Dorsal Raphe Nucleus during a foraging assay. Fluorescence transients were tightly time-locked to reward delivery - here, the appearance of a food pellet. Top: raw DF/F trace. Bottom left: illustration of the optic fibre placement. Bottom right: average activity aligned to pellet delivery time. Peaks in calcium bulk fluorescence from Dorsal Raphe serotoninergic. Shaded areas: standard error of the mean.

**Figure 8.**
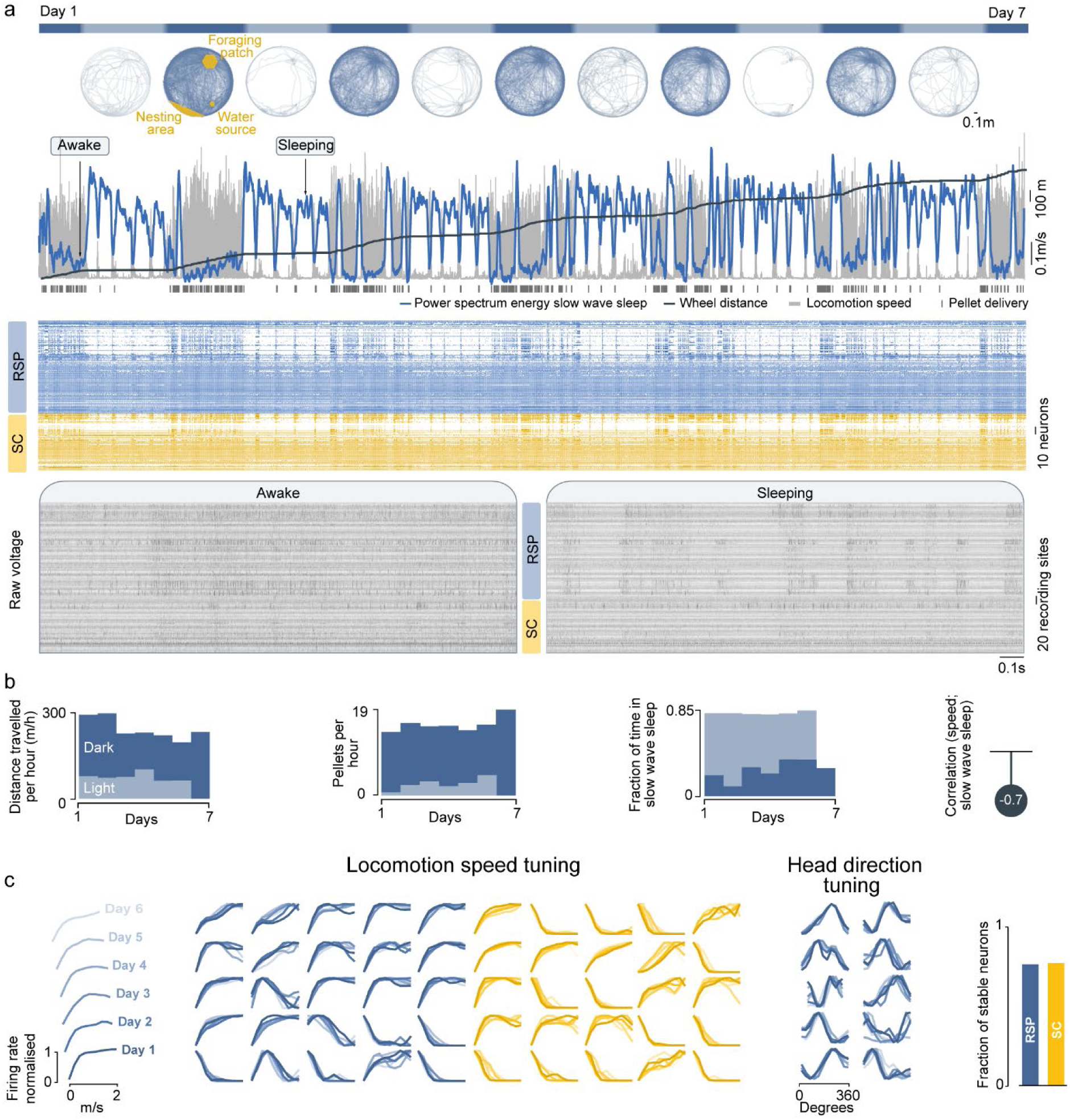
Week-long Neuropixels recording of freely-moving animal during foraging. **a.** Top to bottom: luminance periods (blue bars for low luminance, grey bars for high luminance); trajectories of mouse location in the habitat; locomotion speed time series (light grey area); wheel distance spun time series (dark grey line); pellet delivery time raster (dark grey ticks); 0.5 Hz – 4 Hz (slow wave sleep) cortical power spectrum energy time series (blue line); spike raster plot sorted by brain region (RSP-Retrosplenial cortex, SC-Superior colliculus) and correlation to locomotion speed; raw voltage traces recorded by each of the Neuropixels probe recording site for 1 sec-long awake and sleeping period, showing cortical desynchronised during wakefulness and synchronised activity during sleep. **b.** Left: Global behavioural statistics over days and luminance periods. Right: Correlation value between locomotion speed and the power spectral energy of slow wave sleep. **c.** Left: Example tuning curves to locomotion speed (RSP and SC) and head direction (RSP) for cells whose tuning was classified as stable (see Methods) throughout the entire recording shown in panel **a.** Right: Fraction of RSP and SC neurons with stable tuning.

We recorded the activity of 116 neurons from the retrosplenial cortex (RSP) and 71 neurons from the superior colliculus (SC; Extended Data Fig. 10), two key hubs involved in instinctive navigation and the orchestration of ethological behaviours^3,5,37–39^. These recordings were conducted using Neuropixels 2.0 probes^24^ over the course of a week (Fig. 8a). To track neural identity and analyse neural function over this prolonged time scale, we first spike-sorted the entire duration of the recording in a single pass, then used cross-validation based on unit tuning stability to track units over time (see *Methods*).

We observed that 72% of RSP neurons and 71% of SC neurons showed stable tuning over a week-long period (Fig. 8c). Note that this is a conservative estimate, that does not for instance include neurons showing representational drift^42^. These findings demonstrate the capability of the Aeon platform to maintain precise longitudinal tracking of neuronal activity, enabling comparison of neural dynamics across multiple ethologically relevant conditions and learning.

Aeon’s recording capabilities are not limited to Neuropixels probes. Thanks to the integration of the Onix system, the platform now supports a broad range of technologies—including miniscopes, tetrode-based probes, silicon-probe arrays, tungsten microwires and steel EEG wires^36^. Moreover, Aeon can readily interface with third-party systems that are not Onix or Harp-compatible. For example, a standard non-Harp fibre-photometry setup (see *Methods*) can be synchronised via a TTL pulse sent by Aeon’s Harp camera controller^43^ and detected by the photometry rig’s National Instruments data-acquisition board (Fig. 7d). We used this configuration to record bulk calcium fluorescence from serotonergic neurons in the dorsal raphe nucleus (Extended Data Fig. 9) of mice engaged in a foraging task (Fig. 7e).

This powerful, multimodal integration of recording techniques and behavioural data paves the way for mechanistic insights into how individual neurons, neural populations and neuromodulatory systems orchestrate adaptive and instinctive processes in ethologically-relevant contexts.

## DISCUSSION

Here we describe Aeon, an open-source hardware and software platform to study multi-animal ethological behaviours and their neural substrates over time scales of milliseconds to months in complex and large environments (>2m). Mice housed in an Aeon habitat can express an extensive repertoire of ethological behaviours, including foraging, escaping, nesting, sleeping, drinking, and social interactions. The platform’s modular architecture allows researchers to configure habitats dynamically to fit specific experiment needs, while an integrated suite of sensors and actuators supports deep behavioural quantification. A robust behavioural data acquisition and analysis pipeline, incorporating tools from the Bonsai, Harp, and DataJoint ecosystems, ensures seamless integration of behavioural and neural data, automated data recording, storing, processing, and analysis. Aeon also addresses critical challenges in long-term, large-scale neural recordings. By extending the capabilities of OpenEphys’ ONIX system^36^, Aeon enables continuous, neural recordings in large habitats. By integrating these elements into a fully unified solution, we believe Aeon offers a universal, accessible and standardised system for studying the neural basis of behaviours, as it surpasses currently published solutions, each addressing only a subset of the Aeon requirements outlined in the *Introduction*^18,44–48^.

The Aeon platform has been designed to be fully open source, ensuring accessibility and adaptability for the broader neuroscience community. The platform is supported by a documentation website (https://aeon.swc.ucl.ac.uk/) that provides comprehensive guidance on constructing hardware modules (…/user/aeon_modules/hardware.html#target-hardware), assembling the habitat (…/user/aeon_modules/hardware/habitat.html#target-habitat), designing experiments via Bonsai workflows (…/user/aeon_modules/acquisition.html#target-acquisition-modules), and organising and analysing the resulting datasets (…/user/how_to.html#target-how-to). The core software consists of Bonsai and Python, both freely available. All core packages and modules can be downloaded from the project’s GitHub repositories (Fig. 2b), as well as from NuGet (https://www.nuget.org/) and PyPI (https://pypi.org/). By making all components openly available and documented, we aim to establish Aeon as a standardized ecosystem for neuroscientific experimentation, enabling researchers to develop studies that share common core elements, data formats, and analysis pipelines. Unlike existing data standards such as NWB^29^, Aeon’s format accommodates both structured, trial-based experiment designs and more flexible paradigms, where events are self-initiated by subjects, with behavioural and neural data continuously recorded. Aeon’s modular, paradigm-agnostic design further strengthens its utility by making it makes it broadly applicable to a wide array of experimental paradigms and model organisms ^49–64^. We believe this standardisation of the Aeon ecosystem will not only enhance reproducibility and simplify data sharing, but also facilitate cross-laboratory comparisons, a critical requirement for large-scale collaborative efforts in neuroscience^65^.

A key advancement introduced by the Aeon platform is its ability to conduct continuous, long-term behavioural and neural recordings. This approach offers multiple benefits. It enables researchers to capture slow behavioural dynamics—such as circadian fluctuations, sleep and extended learning processes, which would be impossible in shorter experiments. Additionally, housing mice in a dedicated experiment habitat avoids the confounding stress of removing them from the home cage just before testing. Finally, recording an animal’s extended history— including social hierarchy, food intake, weight, and other behaviours—allows a more thorough characterization of individual differences and enables the interpretation of a given epoch of data to be considered within a broader context. For example, by hosting social groups in long-term experiments, Aeon enabled direct investigation into how social context shapes mouse action policy and learning over extended timescales, demonstrating that mice display more efficient foraging while co-housed. Additionally, the platform’s capacity to record from the same neurons over prolonged periods, while performing high-resolution quantifications of multiple behaviours, greatly expands the scope of research into how the brain organises behavioural sequences and switches between behavioural modules^15,66^. This longitudinal, multi-behavioural perspective also provides a unique opportunity to identify neural populations that are recruited across distinct behavioural contexts—potentially revealing general functions of individual neural circuit that would remain hidden if each behaviour were examined in isolation. Because these recordings span both wakefulness and sleep, the resulting datasets may also constitute a natural window on memory-consolidation processes and how off-line activity could shape subsequent behaviour. This synergy provides the ideal platform for testing computational theories of behaviour and its implementation in naturalistic contexts^32,33,67^, ultimately shedding light on the neural mechanisms that underlie spontaneous and adaptive behaviours.

Alongside its utility for basic neuroscientific research, Aeon’s principles of standardization, modularity, and scalability—combined with its ability to generate detailed, high-dimensional characterizations of both behaviour and neural activity—make it a promising tool for translational research, particularly for high-throughput parallel screening of drug effects and comprehensive phenotyping of mouse models for diseases.

To conclude, by unlocking the possibility of performing system neuroscience experiments in real-world settings, Aeon represents a significant advancement towards gaining mechanistic understanding of how neural circuits implement ecologically relevant computations driving behaviours.

## METHODS

Comprehensive documentation detailing the assembly and operation of Aeon habitats and their components, experiment design within the Aeon ecosystem, as well as data management and analysis workflows, is available on the Aeon project website (https://aeon.swc.ucl.ac.uk/) and associated GitHub repositories (Fig. 2b). Here, we provide explanations and examples to give the readers a general understanding of the methodologies and of the experiments described in the paper. For detailed protocols, step-by-step instructions, and in-depth descriptions, we refer readers to the online resources.

### Animals

All experiments were performed under the UK Animals (Scientific Procedures) Act of 1986 (PPL PP2131611 and PFE9BCE39) following local ethical approval (Sainsbury Wellcome Centre, University College London). Adult Male C57BL/6J wild-type (Charles Rivers) and SERT-Cre mice Jackson Laboratory (stock #014554) were used for experiments. Unless stated otherwise, mice had free access to food and water on a 12:12 h light:dark cycle (ambient temperature 24 °C, humidity 47 relative humidity).

### Aeon habitat construction and functional modules

An Aeon habitat consists of a modular floor made of matte white acrylic (5 mm thick), composed of multiple hexagonal tiles (hexagon side length: 211 mm) supported by a metal framework (Rexroth profiles, dimensions: 60 mm x 30 mm x variable lengths). Each tile fits into a hexagonal slot within an underlying acrylic honeycomb grid, which is designed to accommodate various Aeon hexagonal functional modules (Fig. 1b,d, 2a and Box 1).

#### Empty floor tile module

A flat hexagonal tile used to compose open field areas of the habitat’s floor.

#### Wall corridor and gate modules

In the wall module, a red-tinted see-through acrylic wall (thickness 5 mm, dimensions: 480mm x 350mm) is slotted into an opening in the floor tile and supported from custom made underground acrylic holders. The corridor modules combine 2 parallel walls spaced 4 cm from each other to create a corridor that can be elongated by combining multiple adjacent corridor modules. Some walls present a vertical opening in the middle (width 40mm), that is used by the mouse as a gate to pass through the wall to an adjacent tile (Fig. 1b,d, 2a and 4).

#### Nest module

The nest module is designed to provide a dedicated nesting site and access to water for the mice, while continuously monitoring their weight. The module (Fig. 1b,d, 2a, 3c, and 4) consists of a compartment enclosed by three acrylic walls (dimensions: 210mm × 230mm × 400mm) positioned within the habitat. This compartment is situated above a nesting box, elevated 45mm, which is attached to the balance plate of a scale (Ohaus Navigator NVT Portable Balances NVT4201) for precise weight measurement. A water bottle (Bottle for GM500, TECNIPLAST) is suspended laterally, with only the waterspout protruding inside the nest.

A combination of computer vision and scale readings are used to decide when a weight measurement should be considered. Weight measurements are taken exclusively when a single mouse is present on the scale and the scale readings have been stable for the past 3 s. To prevent baseline drifting, the scale is automatically zeroed frequently, specifically when all mice detected by computer vision away from the nest, exploring the arena.

#### Foraging patch module

The foraging patch module consists of two main systems, each comprising multiple components: the spinning wheel and the pellet delivery system. The foraging wheel is a cylindrical structure measuring 34mm of width and 80mm of diameter, with two distinct sets of 40 pockets engraved on its curved surface: "grip pockets" (dimensions: 4.8mm × 30mm × 1.2mm), which allow the mouse to grasp and rotate the wheel and "pellet pockets" (4.8mm x 4.8mm x 1.2 mm) positioned adjacent to the grip pockets, designed to collect pellets (20mg chocolate-flavoured dustless precision pellets, LBS-biotech) once they have been delivered onto the wheel (Fig. 5a). The wheel’s pivot is supported by two types of ball bearings (Stieber CSK8 and SKF 6082RSL), one of which is unidirectional, ensuring the wheel rotates in only one direction. The wheel’s rotational movement is measured by a magnetic encoder (OEPS7216), mounted parallel to its flat surface, which detects the rotation of a magnet (RS 219-2244) attached to the wheel’s pivotal axis.

Pellets are stored in an underground food hopper (maximum capacity ∼10000 pellets) located in front the wheel under the tile floor and are dispensed onto the wheel via a DC motor-driven spinning disk (Faulhaber 2619S_SR_IE2-16). When the disk rotates, a pellet is delivered through a short chute (chute length: 7 mm) onto the wheel, where it is detected by an infrared beam-break sensor. The beam-break sensor and the motor are controlled by microcontrollers (Raspberry Pi Pico). Data from the beam-break sensor, wheel angle, and motor commands are transmitted and time-stamped by a Harp output expander^26^, which connects the foraging patch module to the behavioural acquisition computer.

Before each experiment, we tested each foraging patch module by coupling its wheel to a motorized gear, spinning at the average mouse spinning speed. We triggered 1,000 pellet deliveries (with a distance threshold of 1 m) and used computer vision alongside IR beam-break signals to detect pellet deliveries and verify proper functioning. If any abnormal behaviour was detected, the foraging patch module was withheld from the experiment and retained for further inspection.

Additionally, to ensure that all wheels required the same effort to spin, we measured the static torque needed to initiate wheel movement before each experiment. The mechanical components on the foraging patch module were then adjusted until each wheel exhibited a torque of 2·10^-3^ N·m. Most mechanical elements composing the foraging patch module can be 3D-printed (material: Stratasys Vero White Plus; printer: Stratasys J750, printer technology: Polyjet).

#### RFID modules

To detect the mouse identity at behaviourally relevant locations within an Aeon habitat, multiple Aeon modules (including the foraging patch module, corridor module and gate module) have been equipped with RFID readers mounted below the tile floor. RFID readers are constituted by a 38mmx40mm receiver antenna (OEPS-3019) connected to a custom-made Harp-compatible PCB board (OEPS-2094). Mice were implanted with RFID chips (Avid FrindChip AVID2328) in the ventral part of their flank, minimising the distance from the antenna and maximising detection efficiency.

#### Custom IR lights

To generate intense and homogeneous infrared illumination within Aeon habitats, we developed custom-made infrared lights. Each light contains 6 meters of infrared LED strips (LEDLightsWorld - DC12V SMD3528-1200-IR), assembled in three 2-meter segments positioned parallel to each other. These LED strips are affixed to a metal case using heat-resistant silicone (RS component, 458-783) and covered by a light diffuser to ensure uniform light distribution. The lamps are suspended to the overhead metal framework at ∼2 m from the habitat floor, with a spacing of one lamp every 60 cm.

### Behaviour-acquisition and ONIX Neuropixels acquisition machines

The high data volume, real-time processing demands, and the need for stable acquisition over extended periods necessitate high-performance hardware for the machines controlling the Aeon platform (Fig. 2c). In table 1, we outline the hardware specifications tested for an Aeon experiment that includes ten FLIR Blackflies cameras (see section High-speed videography in the Aeon environment), four foraging patch modules, one nest module, seven RFID modules, and online SLEAP-based multi-animal tracking, and for the ONIX Neuropixels machine, one Neuropixels 2.0 probe.

**Table 1.** Tested requirements for an acquisition computer to perform an Aeon experiment analogous to the ones described in Fig. 6.

|  |  |
| --- | --- |
| **Processor** | **Intel(R) Core(TM) i9-14900K, 3200 Mhz, 24 Core(s), 32 Logical Processor(s)** |
| **Mother Board** | Z790 AORUS MASTER |
| **Graphic Card** | NVIDIA GeForce RTX 4090a |
| **Operating System drive** | 1TB SAMSUNG 990 PRO M.2, PCIe 4.0 NVMe (up to 7450MB/R, 6900MB/W) |
| **Data storage drive** | 3x4TB SAMSUNG 990 PRO M.2, PCIe 4.0 NVMe (up to 7450MB/R, 6900MB/W) 4th M.2 SSD Drive |
| **Memory** | 64GB to 128GB |
| **Power supply unit** | CORSAIR 1500W HXi SERIES™ MODULAR 80 PLUS® PLATINUM, ULTRA QUIET |
| **Network Card** | Any 10Gbit Ethernet card |

If users wish to customize the hardware configuration to meet their specific requirements, we recommend the following guidelines. Based on our observations, we strongly advise using Intel processors instead of AMD processors, as AMD-based machines consistently demonstrated reduced performance with our specific hardware combination during our tests. For experiments involving multi-animal tracking, the online SLEAP tracking package in Bonsai requires NVIDIA GPUs compatible with CUDA.

For local data storage, we recommend M.2 SSD drives due to their high writing speeds, which are critical for handling the high data acquisition demands generated by multiple high-frame-rate cameras. However, with the use of real-time video compression, standard SSDs may also suffice.

Additionally, a 1 ONIX PCIe Host and a ONIX PCIe Analog and Digital Expansion (breakout board) may be added without significant performance issues to the ONIX Neuropixels acquisition machine ^36^ (Fig. 2c).

### Hardware control and synchronization

Hardware devices within each Aeon habitat are heterogeneous and record different behaviour and neural data modalities, and may also be used to drive, synchronize, and control other equipment. To ensure a common hardware clock reference is in place for all measurements and control signals, we adopted the Harp standard^26^. The Harp clock synchronization protocol allows compliant devices to continuously align their clocks against a reference periodic timing source in hardware, down to a precision of 50 us^26^. All measurements and device commands are hardware timestamped at acquisition time by each device using its synchronized clock. In table 2, we list all Harp devices and peripherals currently used in Aeon platform modules, for ease of reference.

**Table 2.** List of Harp devices used in the Aeon platform.

| **Name** | **Authors** | **Repository** |
| --- | --- | --- |
| Timestamp Generator Gen3 | Open Ephys Production Site | https://github.com/harp-tech/device.timestampgeneratorgen3 |
| Camera Controller Gen2 | Open Ephys Production Site | https://github.com/harp-tech/device.cameracontrollergen2 |
| RFID Reader | Open Ephys Production Site | https://github.com/harp-tech/device.rfidreader |
| Output Expander | Open Ephys Production Site | https://github.com/harp-tech/device.outputexpander |
| Magnetic Encoder | Open Ephys Production Site | https://github.com/harp-tech/peripheral.magneticencoder |
| Sound Card | Hardware and Software Platform, Champalimaud Foundation | https://github.com/harp-tech/device.soundcard |
| Behavior | Hardware and Software Platform, Champalimaud Foundation | https://github.com/harp-tech/device.behavior |

A single Harp Timestamp Generator Gen3 (OEPS, see table2) is used as the authoritative timing source for an Aeon habitat, broadcasting the current time every second to all other devices, either directly from available outputs in the Timestamp Generator, or through daisy-chaining via other Harp devices. All other external devices are directly triggered, or controlled, by this set of synchronized Harp devices, therefore ensuring consistent timing across all experimental equipment.

High-speed videography cameras (described below) are positioned in the habitat and configured for hardware-triggered exposure, using shared trigger lines, either 50 Hz (overhead) or 125 Hz (behaviour hotspots), originating from a single Harp Camera Controller Gen2 (OEPS, see table 2). For every triggered exposure, a Harp-timestamped message is sent to the computer to be matched against the acquired camera frame. This sequence of timestamped exposure events is stored side-by-side with the raw video sequence.

Foraging patch modules are controlled directly using Harp Output Expander devices (OEPS, see table 2), which include a magnetic encoder peripheral (OEPS, see table 2) used to track angular wheel movement, and various digital I/O lines to control and monitor pellet delivery. Both continuous wheel data and digital I/O events and commands, such as pellet delivery triggers, and pellet delivery beam breaks, are logged with hardware timestamps.

Harp RFID Reader devices (OEPS, see table 2) can be positioned across the arena to detect 125 kHz RFID microchip implants on multiple animals. In the specific case of the RFID Reader module, the Harp synchronization signal is sent via the host computer to minimize device footprint and cable requirements. The readout latency of the RFID tags is very high (on the order of 300 ms) relative to the clock precision (50 us) and device latency (1 ms), making timing deviations negligible for identification purposes. For precise real-time positioning the identification data is combined with other tracking sources.

For continuous monitoring of animal weight, we used a weighing scale placed directly underneath the nest module, and connected to the acquisition system via USB. As the weighing scale itself is not a Harp device, and the readout latency is high (on the order of 50 ms) relative to the Harp device communication latency (1 ms), we synchronised weighing measurements in software against a Harp hardware timestamp source streaming at 500 Hz.

Auditory stimulus can be delivered using either Harp SoundCard^26^ (OEPS, see table 2) or a Harp Behavior (OEPS, see table 2). The Harp SoundCard allows playback of audio waveforms up to 192 kHz with a linear frequency response^26^ when coupled with the Harp Audio Amplifier (OEPS, see table 2). The Harp Behavior device can be used to drive PWM signals either for auditory (coupled directly to a piezo buzzer) or optogenetic stimulation (coupled to an LED source) with frequencies up to 10 kHz.

Other software-only data streams of interest include habitat metadata and transition events, such as animals entering or leaving the habitat, manually restocking the pellet dispenser, environment lighting control, or changes in the environment logic rules and parameters. These “soft” events are currently timestamped in software, typically against the overhead camera exposure timestamps, as their timing precision is on the order close to a second (as they depend either on human manual input or are slow changes). Still, there is an advantage in having them timestamped against a known time reference as they can be used during data analysis to split up and filter the data into different experimental conditions.

### High-speed videography in the Aeon environment

The Aeon platform supports a broad range of camera models (e.g., Teledyne FLIR and Basler). In the habitats used for this paper, we used FLIR USB3.0 Blackfly-S cameras (Model BFS-U3-16S2M-CS, 1440×1080 pixels). In a typical experiment we would use 6 top view cameras (50 Hz), and 3 side-view cameras zoomed in on the foraging wheel (125 Hz). The acquisition of every frame was triggered by TTL pulses generated by a Harp Camera Controller Gen2 device (described above, table 2).

### Experiment control and design in Aeon habitats

Experiment control, data acquisition and logging in Aeon habitats are entirely implemented using the Bonsai programming language^28^. Bonsai supports many core and third-party packages for integrating with a variety of commonly used software and hardware. Notably, the Bonsai.Harp package enables direct communication with all Aeon Harp devices using the standard Harp binary communication protocol. Additionally, we developed several Aeon-specific Bonsai packages, including Aeon.Acquisition, Aeon.Database, Aeon.Environment, Aeon.Foraging, Aeon.Video, and Aeon.Vision, among others. These packages support Aeon’s standardized data logging format, enable communication with MySQL databases, provide webhook connectors to broadcast alert messages during an experiment, implement persistent storage of configuration state to allow recovery in the event of an unexpected crash, interface with a variety of devices including the foraging and camera modules, and integrate SLEAP for real-time identity tracking and pose estimation, among many other diagnostic, control, and data acquisition features. All packages have been published open source on the NuGet gallery (https://www.nuget.org/) and can be readily installed into any Bonsai experiment.

The Bonsai workflow files that run on the behavioural acquisition computer during an experiment heavily incorporate components from the Aeon-specific Bonsai packages mentioned above, and share some infrastructure design patterns which are recurrent in our experiments. All experiment workflows for experiments described in this work live in a single code repository, aeon_experiments, in which the ‘main’ branch serves as an experiment template. Every branch in this repository is rooted in this ‘main’ branch, and contains that experiment’s specific logic for habitat control, dynamic habitat updates, device calibration, data chunking, data transfer, and state recovery. As an example, in the ‘social’ branch which governs the social foraging assay described in Fig. 6 and Extended data Fig. 7, Bonsai code controls the light cycles, updates patch properties according to parametrized rules, calibrates the overhead cameras, transfers data at the end of every hour-long chunk to network storage, and saves and reloads experiment state following a system crash and restart. Experiment workflows are structured in top-level group nodes that encapsulate code for general experiment control, experiment and device metadata definitions, data logging of specified devices, dynamic triggering of habitat updates, GUI-based habitat monitoring and updating, and real-time system alerts.

The rules governing high-level environment logic are themselves implemented in Bonsai and parameterized by environment configuration files (“environment.yml”) to allow experimenters to specify long-term environment conditions such as the type and the dynamics of each foraging patch (Extended Data Fig. 4 and Extended Data Fig.7c-e). Each experiment may define its own environment definition language with specific parameter sets by means of a JSON schema, which is used to validate and load all configuration files. Environment rules can be changed at any time during the experiment, and therefore the environment configuration files themselves are timestamped and saved as part of the data whenever they are loaded, so that the experimenter can know which rules were active in the habitat at any given time.

### Data logging, preprocessing and analysis in the Aeon platform

Each data stream has a distinct identifier within its device, which is augmented with the chunk UTC suffix to uniquely name data files even when recording over daylight saving periods. All data from a device is stored in a folder with the device name, containing all collected device data stream files, and a data “epoch” is defined as a continuous period of acquisition in which the system is turned on and collecting data uninterrupted. All data from an acquisition epoch is stored in a folder containing all devices and data streams. An experiment may be composed of multiple epoch folders if the system experiences (and recovers) from a critical failure or requires human maintenance demanding for a system interruption at any point.

### DevOps practices in the Aeon platform

In order to enforce best development practice and standardise data format and architecture, the Aeon platform adopted a set of DevOps practices, which provide streamlined project management and practical integration and deployment processes. We make extensive use of GitHub for all project and task management. All development efforts are traceable though a repository’s milestones and issues. To ensure software reliability, GitHub Actions automate continuous integration (CI) testing across all core Aeon repositories. Deployment to experiment acquisition computers, however, remains a manual process, ensuring experimenters retain full control over system updates. To allow experimenters to refine experiment setups without needlessly storing incomplete datasets, deployment practices enforce commit tagging and repository cleanliness to differentiate validated experiment data and test data. Acquired data is considered valid experiment data and stored in the final raw data location only if the latest commit in an experiment branch of the aeon_experiments repository on an acquisition computer is tagged and the repository is in a clean state. Conversely, untagged or uncommitted changes in the repository are treated as test data.

### Classification of behaviours

#### Sleeping

Sleeping episodes were extracted from the video frame recorded by a zoomed in camera pointing down at the nest (Fig. 4). A sleeping bout was defined as a continuous period of complete immobility lasting for at least 2 minutes. Gaps smaller than 100 ms were ignored.

#### Drinking

Drinking episodes were extracted from the video frame recorded by a zoomed in camera pointing down at the nest (Fig. 4). Video frames were first analysed using a SLEAP model that extracted x,y coordinates of the animal’s snout tip and body centre. A drinking episode was defined as the time period during which the mouse snout was within 30 pixels from the tip of the water bottle spout for at least 0.5 s. If the mouse snout or body centre was detected between the water bottle spout and the nest wall where the bottle was mounted, the frame was excluded.

#### Escaping

Escapes were triggered following the procedure described in Vale et al 2017^68^. Threatening stimulus was a 15kHz tone (sound pressure at the arena floor 80dB) delivered using an overhead ultrasound speaker (L400, Petterson) controlled and time-aligned by a Harp soundcard and a Harp behavioural board. Onset of escape was determined by visual inspection of the video recordings and considered as either the onset of a head-rotation movement towards safety or an acceleration (whichever occurred first) after the mouse detected the stimulus^5^.

#### Feeding

Under our experiment conditions, mice do not harvest pellets and virtually always consume them as soon as they are delivered. We therefore used pellet delivery time as a proxy for putative feeding episode time.

#### Exploration

Exploration epochs were defined as time periods during which the mouse was outside the nest.

#### Bouts of aggression

Bouts of aggression were identified using a multi-step process based on SLEAP pose tracking. Initially, frames were marked as potential aggressive events when the centroids of the two mice were within close proximity (approximately 3.7 cm) and when their tracked skeletons indicated implausible body parts—such as exaggerated distances between body parts on a mouse’s head or unnatural spacing along their spine—all of which commonly result from rapid, chaotic movements and overlapping postures typical of aggressive interactions. We also incorporated frames with complete tracking failures (empty frames) when they occurred while mice were known to be in close proximity based on adjacent frames. Candidate frames occurring closely together in time were grouped into preliminary bouts.

Individual mouse speeds were then analysed within these preliminary bouts, as aggressive interactions typically involve rapid and pronounced movement. This step required careful verification of tracked mouse identities to accurately calculate each mouse’s speed. Aggression was confirmed only if either one mouse exceeded 20 cm/s or both mice exceeded 15 cm/s on average, distinguishing genuine bouts of aggression from other close but less active interactions. Finally, only bouts lasting at least one second were retained to minimize false positives from brief, spurious detections.

#### Social encounters and retreating

Social encounters and retreats were detected by analysing mice positions and orientations using SLEAP pose tracking. Frames were initially flagged as candidate events when the mice were facing each other within close proximity (approximately 9 cm). To confirm that the mice were truly in head-to-head confrontations, we required opposite orientations within a 45-degree tolerance and implemented spatial constraints that distinguished these interactions from side-by-side or back-to-back configurations. Consecutive frames meeting these criteria were grouped into candidate encounters when they occurred within a brief time interval.

Each candidate encounter was then screened for tracking errors, particularly skeleton flipping cases where a mouse’s tail might be misidentified as its head, resulting in false detections. Events containing such errors, indicated by inconsistent orientations between frames, were discarded from further analysis.

For each remaining candidate encounter, we established a 1-second window following the initial interaction to search for a retreat event, which would confirm a complete social encounter-retreat sequence. Within this window, we looked for frames showing that one mouse had turned around (i.e., they now shared a similar orientation, within a 45-degree tolerance). To robustly confirm these tube test end frames, we enforced a movement threshold requiring the loser’s average speed to exceed 2 pixels/s to exclude stationary behaviours that also involve orientation changes like grooming.

#### Foraging bouts

Foraging bouts were defined per mouse as time intervals in which the mouse initiated at least three pellet deliveries and rotated the patch wheel by at least one full turn within a 60-second period.

### Foraging patch preference experiment and quantification

To estimate patch preference mice were located in a habitat with two foraging patch modules, with the same distance threshold. After the mouse have sampled both patches and collected a total of about 20 pellets, the threshold of the patch with the highest pellet count was increased to a new value (see Fig. 5c,d). Patch preference was computed as the difference between the distance spun from the switch time over their sum.

### Classification of behavioural syllables using Hidden Markov Models

#### Hidden Markov Models (HMMs)

HMMs were trained using six factors as inputs: mouse kinematics (mouse centroid speed and acceleration magnitude) and four pairwise distances between mouse body parts estimated using SLEAP tracking (Extended Data Fig. 5). The most relevant pairwise distances were selected by performing Singular Value Decomposition (SVD) on a standardized distance matrix and identifying the features which capture the most variance in the data. These features were the distances between head and spine 3, spine 1 and spine 3, left ear and spine 3 and right ear and spine 3, that captured both the mouse body length and body curvature.

HMM was implemented using the SSM library^69^, specifying a Gaussian emission distribution for the observations. The model was initialized with a predetermined number (*m*) of hidden states and the dimensionality (*k* = 6) of the observation space X = *x*_1_, *x*_2_, ⋯, *x*_*T*_. Each observation at time *n* is represented by a vector *x*_*n*_, and the corresponding latent state is denoted as *z*_*n*_. The Expectation-Maximization (EM) algorithm was used to iteratively estimate the model parameters *θ* to maximize the likelihood of the example observed data:

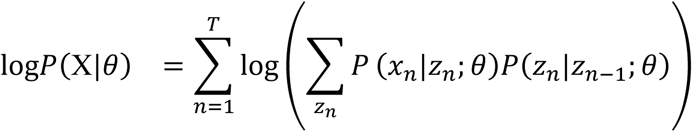

The fitting process was conducted with over 50 iterations. The optimal number of latent states was determined by comparing the average cross-validated log-likelihood per point across different models: each model was fitted using a 24-hour sample dataset; the Viterbi algorithm was then applied to infer the most likely sequence of hidden states for observations from the second full day of the session. The average log-likelihood per point was calculated by dividing the overall log likelihood of this inference by the total number of observations. Models with higher log-likelihood values were preferred, as they indicated better fit to the data.

The number of states was determined as *m* = 10. After training the model on active chunk data, the Viterbi algorithm inferred the most likely sequence of hidden states for the full session. Parameters for each of the six observation factors corresponding to each state were obtained, including the mean and covariance matrix for the Gaussian emission distribution of each factor. States were sorted in ascending order based on their mean speed parameter. The transition matrix was correspondingly reordered to match the sorted states. The Kullback-Leibler (K-L) divergence between two states was calculated as follows:

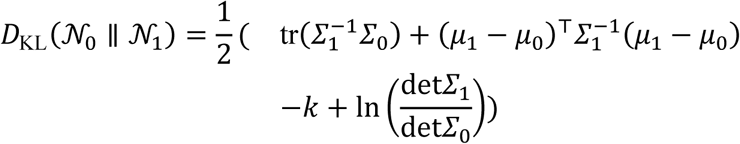

where *μ*_0_ and *Σ*_0_ are the mean vector and covariance matrix of the first state 𝒩_0_, *μ*_1_ and *Σ*_1_ are the mean vector and covariance matrix of the second state 𝒩_1_, and *k* is the dimensionality of the state (*k* = 6).

To correct potential tracking errors, the mouse (i.e., position, velocity and acceleration) were inferred using a discrete Wiener process acceleration model and a Kalman filter algorithm^70^.

#### Statistical modelling to predict event

The behavioral states were converted into regressors by calculating their occupancy probabilities over fixed time intervals. Specifically, data was sampled every 10 seconds *Δt* = 10 to align with the regression model’s time step. In the model, each behavioral state *i* at time *t*, *S*_*i*_(*t*) represents the occupancy probability, and *s*_*i*_(*t*) represents the number of occurrences of state *i* in the past 10 seconds. The occupancy probability is calculated as:

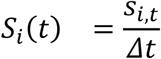

expressing the proportion of time that state *i* was observed within each 10-second interval.

A kernel density estimation (KDE) method was applied to quantify the temporal intensity of behavioral events. The intensity function *λ*(*t*) was calculated using a Gaussian kernel with a bandwidth scaling parameter *h* = 0.001:

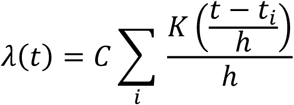

For the *i*_*th*_ event occurring at time point *t*_*i*_, we applied a Gaussian kernel function 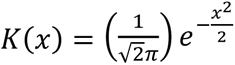 to estimate its temporal contribution to the overall intensity. These individual kernel contributions were then aggregated and normalized by a constant *C* to produce a smooth, continuous intensity function across the observation period. The resulting intensity estimates were normalized by dividing by the maximum intensity value, producing a standardized intensity function ranging from 0 to 1. To incorporate these intensity values into our subsequent analyses, we mapped the smoothed intensities back to the original temporal resolution of our dataset using nearest-neighbor interpolation.

A Gaussian-Generalized Linear Model (GLM) was applied to the active chunk of the data to predict the intensity function:

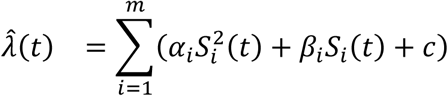

where *α*_1,⋯,*m*_ and *β*_1,⋯,*m*_were the coefficients for each state, and *c* was the constant term. Models were evaluated based on the correlation between their prediction and the event intensity function. After cross-validation, the model with the highest correlation coefficient was selected.

To validate the model’s performance and establish statistical significance, a permutation test approach was implemented. The temporal sequence of state occupancy probabilities was randomly shuffled while maintaining the probability distributions at each timepoint. Using these shuffled input, new predictions were generated and their correlations with the true intensity function were calculated. The 95% confidence interval of these correlation coefficients was computed using a standard normal distribution approximation, with the upper boundary (*μ* + 1.96*σ*) serving as the ‘by chance’ threshold.

#### Sequence variability analysis

We analyzed the variability of the behavioural sequences of three foraging related events: visit initiation, visit termination, and pellet delivery. A patch visit was defined as the mouse staying in a foraging patch for > 5 seconds while moving the wheel. Visit start and end were determined by wheel movement, detected by the wheel motion sensor (Fig. 5a). Movements separated by less than 5 seconds were considered parts of the same visit. Visit initiation was marked by the first wheel movement after a period of inactivity. Visit termination was recorded when wheel movement stopped, and the mouse left the foraging patch.

A time point-wise state sequence matrix for each foraging-related event was constructed to summarize behavioral states across multiple events. Each row represented the state sequence of an event, aligned to the trigger of event onset. For each time point, the most frequent behavioral state across all events was identified as the typical state, forming the dominant state sequence. For each foraging event scenario, we defined early events as the first sixth of the sequence and late events as the final sixth of the sequence. The similarity between individual event state sequences and the dominant sequence was quantified using Dynamic Time Warping (DTW) distance.

To compute the similarity of two state sequences (*Seq*_1_, *Seq*_2_) of lengths *l*_1_ and *l*_2_, a DTW cost matrix *D* of size (*l*_1_ + 1) × (*l*_2_ + 1) was constructed. The first row *D*[0] and column *D*[: 0] of the matrix were set to infinity, except for the first element *D*[0,0] being initialized to 0. For each cell *D*_*i*,*j*_(1 ≤ *i* ≤ *l*_1_, 1 ≤ *j* ≤ *l*_2_), its value was computed as:

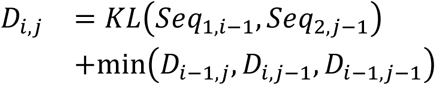

where *KL*( ) is the K-L divergence between states at *Seq*_1,*i*−1_and *Seq*_2,*j*−1_. Ultimately, the DTW distance between sequences was given by 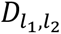, with smaller distance indicating greater similarity with the dominant sequence of the event.

### RFID implantation and tail tattooing

Before the start of any multi-animal experiment, mice were tattooed and implanted with an RFID chip. Mice were anesthetized using isoflurane (3–5% in oxygen). Lubricating ointment was applied to the eyes, Carprofen (5 mg kg^−1^) was administered subcutaneously for analgesia. RFID chips were implanted in the mouse flak by using a pre-loaded single-use RFID syringe (Avid FrindChip AVID2328), with the chip placement verified with an RFID scanner (Smoostart B091FHNL1Y). For tail tattooing, the fur was carefully removed from the tail with a sterile scalpel, the skin cleaned with alcohol, and Vaseline was applied. Using a tattoo machine (Vettech - VTS Tattoo Kit), loaded with 3-point fine needles (Vettech - ID002) and black ink (Color Co - Dynamic TBK ink), one or more bands were tattooed on the tail (band length approximately 1 cm, spacing between bands approximately 0.5 cm). After the procedure, Vaseline was reapplied daily for 5 days to aid healing.

### SLEAP-based identity tracking

Aeon’s tracking pipeline relied on two types of SLEAP models: one for pose tracking and one for identity tracking. Below, we outline the details of the pipeline used for our social foraging experiments, which could be readily adapted to different setups, such as varying camera views, frame rates, resolutions, and numbers of body parts to track, among other factors. Software and documentation to implement the aeon SLEAP pipeline can be found in the aeon_acquisition, aeon_experiments and aeon_sleap_processing github repositories.

The SLEAP pose model tracked eight body parts along each mouse (Fig. 6a and Extended Data Fig. 5 and 6) and was designed to be reusable across experiments with similar arena setups. This model was trained on 650 frames from seven different one-hour full-field videos, each recording up to 3 mice, yielding a total of 1621 labelled mouse skeletons. These videos were selected to capture variations that could affect SLEAP’s predictions, such as differences in lighting, arena surroundings, camera angles, etc.

In contrast, the SLEAP identity models were unique to each Aeon experiment and the specific subjects that were involved. Below is an overview of the tracking pipeline implemented at the start of each new experiment for training the identity model and running inference, ultimately generating combined pose-identity data for the entirety of the experiment.

#### Training data collection

Each Aeon experiment’s SLEAP identity model was trained on composite videos constructed from frames captured by four overhead quadrant cameras, each covering roughly a quarter of the arena with slight overlaps between their fields of view. Although these cameras recorded different sections of the environment, the symmetry of the arena allowed for training a single unified model effective across all four quadrant views.

One-hour videos of each animal alone and a one-hour video of both animals together were acquired prior to the social phase of the experiment. To ensure a diverse training dataset, we verified that the animals explored the full extent of the habitat, covering all spatial regions. For each video frame, we identified which quadrant camera provided the most central view of each animal. These frames were then sequentially combined to create composite quadrant camera videos.

For instance, if both animals were best captured by the same quadrant camera, only that single frame was included. However, if they occupied different areas of the arena, two separate frames – each from a different quadrant camera – were selected for that time point and arranged consecutively in the composite video. A corresponding CSV file was generated alongside each composite video to log these selection decisions.

#### Labelling

For the solo training session composite videos, we implemented automatic ID labelling with the position marker placed at each animal’s centre of mass. For the social training session, ID labelling required manual annotation using the SLEAP GUI. We labelled approximately 1000 frames per session, resulting in a total dataset of around 3000 frames for two mice. This corresponded to roughly 2000 labels per mouse: 1000 from the solo session and 1000 from the social session.

#### Training

To train an optimal SLEAP model, we conducted sweeps over various model hyperparameters using the optimization framework Optuna^71^. The first Optuna trial was initialized (“enqueued”) with parameters known to generally produce reliable results for our setup, followed by 100 additional trials. Each model was only partially trained, stopping after 75 epochs to reduce computational time, and was then evaluated on a test dataset, with model performance serving as the objective function for Optuna’s optimization process.

The evaluation metric was a composite measure that combined identity accuracy and instance detection. We included the instance detection component because relying solely on identity accuracy could lead Optuna to produce low-quality SLEAP models that detected very few mice but happened to identify them correctly by chance, thus artificially boosting their accuracy scores. We defined identity accuracy as the proportion of correctly identified instances among all detected instances:

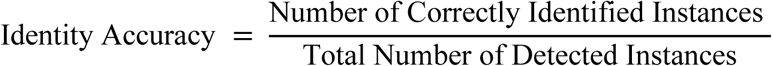

We quantified detection performance using the F1 score:

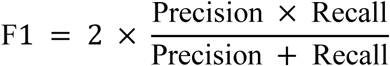

Finally, we combined these two measures via their harmonic mean, which was chosen as it strongly penalizes low values in either metric, ensuring that optimized models performed well in both identification and detection tasks:

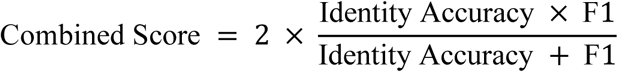

Based on the Optuna results, a complete SLEAP model was then trained using the optimised parameter values and in turn evaluated on unseen, automatically labelled single animal videos to validate performance.

#### Inference

Following training, we deployed the final SLEAP identity model on all four quadrant cameras while running the SLEAP pose model in parallel on the full-field view camera. Both models could perform inference in real time at 50 Hz. To produce a unified pose-identity dataset, we followed a three-step process: first, we selected the highest-likelihood identity prediction for each mouse from across the quadrant cameras. Second, we projected these identities onto the full-camera view using homographies trained with the ORB (Oriented FAST and Rotated BRIEF) algorithm. Lastly, we matched each pose with its corresponding identity using the Hungarian algorithm, a classic approach to the assignment problem that finds the optimal one-to-one pairing by minimizing the sum of distances between predictions (Extended Data Fig. 6). To exploit the redundant coverage afforded by the overlapping full-field and quadrant views, we also run a SLEAP identity model on the full-field camera in parallel; its predictions are seamlessly integrated to fill potential gaps from the quadrant feeds, increasing the identity dataset continuity.

#### Temporal-based error correction

We developed a temporal-based error correction algorithm to correct occasional identity misassignments in the tracking data. The algorithm works in three steps: first, for each frame containing both animals, it calculates two possible identity assignments (maintaining current identities vs. swapping them) and chooses whichever minimizes the total distance each animal would need to move from the previous frame. Second, the algorithm processes data in segments, deliberately excluding periods when animals are within 20 cm of each other since identity swaps are difficult to detect reliably when animals are in close proximity. Third, within each continuous segment, it applies majority voting across all frame-by-frame decisions to account for potential erroneous identity assignment in the first frame of a segment.

### SLEAP-based and RFID-based identity tracking evaluation

To validate the accuracy of the identity tracking pipeline when deployed and assess how it evolves throughout the days of the experiments, we leveraged the arena’s numerous RFID readers as ground truth.

We began by quantifying the range, latency, and detection probabilities of RFID readers. RFID reader detection ranges were determined by matching RFID detection timestamps with concurrent SLEAP pose data within a 10ms temporal tolerance. For each RFID detection, we calculated the Euclidean distance between the mouse’s tracked position and the known RFID reader coordinates and the mean of these distances defined each reader’s effective detection range.

We identified discrete "visits" to RFID readers by defining periods when mice remained within the reader’s mean detection range for ≥1 second. For each visit, we measured the RFID detection latency as the time between the mouse entering the detection zone and the first RFID detection during that visit. Detection probability was calculated as the proportion of visits that resulted in at least one successful RFID detection.

Given the RFID readers’ demonstrated spatial and temporal precision, we used their detections as ground truth for SLEAP identity tracking validation. Importantly, we restricted our analysis to RFID reader “visits” lasting ≤2 minutes to exclude periods when mice were sleeping directly on or near RFID readers, which would generate excessive detections and artificially bias accuracy metrics without reflecting the system’s performance during active behaviours.

Identity classification accuracy was computed daily as the proportion of correctly identified animals among all SLEAP identity predictions that could be matched to concurrent RFID. We calculated this metric separately for each day of the social phase to assess temporal stability of the tracking system throughout the experiment duration.

### Manual and automated tube test

The manual tube test procedure was adapted from Fan et. al 2019^35^. Briefly, after a habituation period of at least 5 minutes during which both mice were allowed to explore and walk through a narrow clear Perspex tube, the testing phase began. Two mice were simultaneously placed at opposite ends of the tube and allowed to walk inside. The loser was defined as the mouse that first retreated and exited the tube. With the tube ends randomized for each trial, the test was repeated a minimum of 15 times and continued until the win-rate for the dominant mouse was determined to be statistically significant (p < 0.05) under a binomial distribution with mean = 0.5.

For the automated tube test, we considered all the “social encounter and retreating” events automatically scored as described in the section *Classification of behaviours*. The loser was classified as the mouse that first turned 135 degrees in either direction from the start of the encounter, as determined by nose pose tracking.

### Neuropixels probe recordings

#### Surgical procedure

Neuropixels 2.0 probe was chronically implanted by cementing it to the mouse skull as described in Campagner et al 2023^3^ and here https://www.youtube.com/watch?v=dO6dMSjuK1g.

#### Histology

Perfusion and imaging using serial section 2-photon tomography was done as detailed in Campagner et al 2023^3^.

#### Neuropixels commutation-translation module

The Neuropixels commutation-translation module encompasses two systems: a linear slider that allows to compensate for the mouse 2D location in the arena and the ONIX commutator (Newman et al 2024) that tracks mouse rotations.

The linear slider is composed by a micro linear actuator (Igus, ZLW-1040-02-100-L-600) operated by a Brushless DC-servo motor (4490-BS, Faulhaber). A Bonsai workflow tracks online the 2D location of the implanted mouse in an Aeon habitat from the full field top view camera images (50 Hz), and for each frame it calculates the desired cable length based on the radial distance between the mouse position in the habitat and the vertical centre of the tether guide to the arena floor. From this desired cable length, the position of the linear stage along the rail is computed and sent to the motor itself via a Raspberry Pi Pico microcontroller implementing the Aeon Linear Drive Harp device (Fig. 2b) via the MicroHarp MicroPython framework (https://github.com/SainsburyWellcomeCentre/microharp). A custom acrylic cable guide is mounted under the liner actuator and prevents the formation of loop in the Neuropixels tethered when the commutator translates along the linear slider. All cables connected to movable part have been enclosed in a cable chain (Igus 7, part number: 07.40.028.0) to avoid loop formation and ensuring longevity across weeks long experiment of continuous movement.

#### Recording and analysis of single units’ activity in freely moving animals using Neuropixels probe

Extracellular recorded voltage was acquired at a sampling rate of 30 kHz using Neuropixels 2.0 probes connected to the ONIX system, controlled via the OpenEphys.Onix1 package. The Harp clock signal is acquired directly by the ONIX system and timestamped in the same clock as other physiology data. All data, including the Harp clock timestamps, IMU and Neuropixels 2.0 channel voltages were chunked into one-hour segments. The size of each raw segment is computed to have the exact number of samples corresponding to one hour acquisition at the intended sampling frequency. In the case of the ONIX recordings we use the Harp clock timestamp stream to synchronize the data with other data streams automatically as an intermediate step during analysis. Spike sorting was performed using Kilosort 2.5 as described in Campagner et al., 2023, with channels divided into three equally sized groups contiguous along the shank. Each group was spike-sorted over the full duration of the experiment (∼1 week).

Tuning curves for locomotion velocity (SC and RSP) and heading direction (RSP) were computed as described in Campagner et al., 2023^3^, using 10 bins for locomotion speed and 12 bins for heading direction, after that the data have been chunked in time into six segments. Heading direction was defined as the direction of the mouse’s velocity vector calculated between consecutive frames. Stability was assessed by generating a null distribution of tuning curve correlations, obtained by calculating the mean pairwise correlation between tuning curves from a shuffled dataset (100 shifting) across different time periods. The stability criterion required the real mean correlation to exceed the 95^th^ percentile of the null distribution.

### Fibre Photometry

#### General surgical procedures

Stereotactic surgical procedures were performed under anaesthesia (1-2% isofluorane, 1L/min) and analgesia was given subcutaneously (Metacam 25µl/10g). Viral vectors were administered using glass pulled pipettes (3.5” Drummond #3-000-203-G/X) with tips clipped to a diameter of ∼20µm. To target Dorsal Raphe we used coordinates 0mm ML, −4.3mm AP, −3.5mm DV relative to bregma and approached this with an anteroposterior angle of 25-30 degrees to avoid the transverse sinus. Injections were performed (5nl per injection cycle, 1nl/s and 10s wait between cycles) using a motorised injector (Nanoject 3, Drummond) and 5-10 minutes were given before retracting the glass pipette to reduce excessive spreading of the injected volume.

For photometry recordings 200nl of flex-GCaMP6f (Penn Vector Core, LOT Number: CS0204, AAV1.Syn.Flex.GCaMP6f.WPRE.SV40, titre: 2.88E+13) was injected into the DR of heterozygous Sert-Cre mice followed by implantation of a fibre optic cannula (Doric Lenses 0.57NA, 400µm diameter) into the DR secured using UV curing dental cement (3M RelyX Unicem 2 Automix) and reinforced with non-UV curing dental cement (Super-Bond).

#### Histology

See Neuropixels probe recordings section.

#### Data acquisition and preprocessing

Photometry data were acquired using custom scripts in Python (https://github.com/stephenlenzi/daq_acquisition) a NIDAQ (National Instruments, USB-6341) together with a Fluorescence Minicube (Doric Lenses, iFMC6_AE_S) and pigtailed rotary joint (Doric Lenses, FRJ_1×1_PT). All other optical acquisition components had a numeric aperture of at least 0.57, and stimulation components 0.22 (Doric lenses). We followed a protocol for acquiring signal and background channels from GCaMP6f expressing neurons by stimulating at calcium sensitive (465nm) and insensitive (405nm) wavelengths that allows ratiometric measurements, bleaching and artifact correction with a single fluorophore^72^. LED output power was matched for the two channels and was amplitude modulated with a sine wave (peak amplitude 0.2 µW) at different frequencies for each channel (211Hz and 531 Hz, respectively) to enable source separation after data acquisition. Photometry signals were preprocessed and demodulated using custom Python code (https://github.com/stephenlenzi/daq_loader).

## AUTHOR CONTRIBUTIONS

Conceptualization: D.C., G.L., J.B. and SGEN group; Software: G.L., J.B., T.T.N, A.A., C.H.L, T.R., B.C., A.E., A.G.P., J.A., M.M., D.C., SGEN group; Hardware: SGEN group, L.C., A.A., D.C., F.J.C., T.R., G.L. J.B.; Data Analysis: J.B., A.G.P., D.C., Z.L., J.R., O.F.; Experiments: D.C., J.B., L.C., S.C.L, J.D.S.R.; Writing: D.C., J.B., G.L., A.G.P.; Documentation and Website: C.H.L., L.C., T.R., G.L., J.B., S.N., D.C, A.G.P.

## CODE AVAILABILITY

https://aeon.swc.ucl.ac.uk/user/index.html,

https://github.com/SainsburyWellcomeCentre/aeon_api,

https://github.com/SainsburyWellcomeCentre/aeon_mecha,

https://github.com/SainsburyWellcomeCentre/aeon_experiments,

https://github.com/SainsburyWellcomeCentre/aeon_acquisition,

https://aeon.swc.ucl.ac.uk/user/how_to.html#target-how-to

https://github.com/SainsburyWellcomeCentre/aeon_scratchpad/tree/aeon_platform_paper_biorxiv/aeon_analysis/aeon_platform_paper

## DATA AVAILABILITY

https://aeon.swc.ucl.ac.uk/user/sample_datasets.html

## AKNOWLEDGEMENT

This work was funded by Sainsbury Wellcome Centre Core Grant from the Gatsby Charitable Foundation and Wellcome (GAT3755 and 219627/Z/19/Z), Gatsby Unit/SWC Joint Research Fellowship in Neuroscience, the SWC PhD Programme, BBSRC Research grant - BB/W019132/1, 2023-2024 Shenoy Undergraduate Research Fellowship in Neuroscience - NC-SURFiN-00003556. The Aeon project was the result of a large collaborative effort. The full contributors list, indicated in the author section as the SGEN group, can be found here (https://aeon.swc.ucl.ac.uk/about/people.html).

