## Extended Data for "Aeon: an open-source platform to study the neural basis of ethological behaviours over naturalistic timescales"

### EXTENDED DATA FIGURES

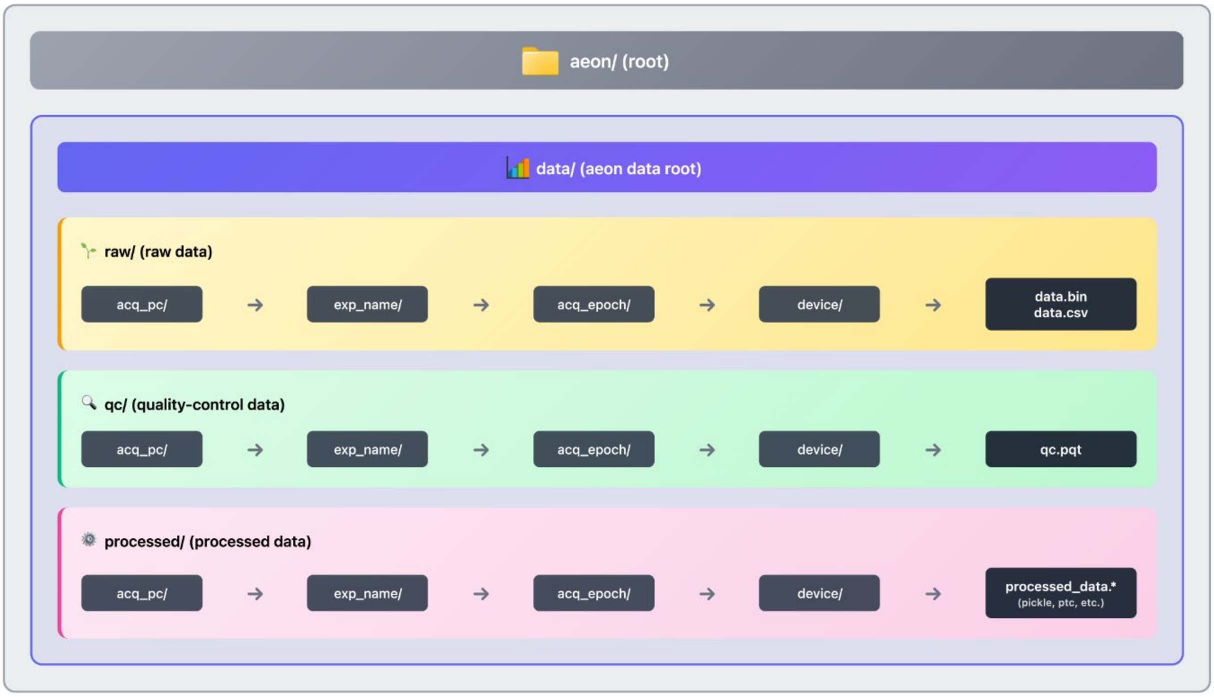


**Extended Data Figure 1 – Directory structure underlying the Aeon data format**

A schematic overview of the default, automated file directory structure of acquired data from Aeon experiments. After an initial root path is specified, all experiment data lives in an associated data root path (‘data/’). There are three classes of data, ‘raw’ (‘aeon/data/raw/’), ‘quality-control’ (‘aeon/data/qc/’), and ‘processed’ (‘aeon/data/processed/’). Raw data is data acquired and recorded directly from streams in a Bonsai experiment workflow. The data acquisition PC (‘acq_pc’) that the Bonsai experiment workflow runs on is the only PC that has write-access to this experiment; i.e. all raw data for a given experiment (in ‘exp_name/’) is write-protected to the PC that acquired this data, guaranteeing provenance and integrity of raw data. QC data is generated per stream via online quality-control processes that run on acquired raw data – e.g. the times of dropped camera frames, and the times a pellet delivery attempt was not followed by a pellet beambreak. These are stored in parquet table file format, allowing for quick and intuitive access indexed by time and event. Lastly, processed data is generated post hoc—for example, we can train an improved identity and pose estimation model after the experiment has concluded and re-run it on the original raw camera data to enhance the results of the initial online estimation.

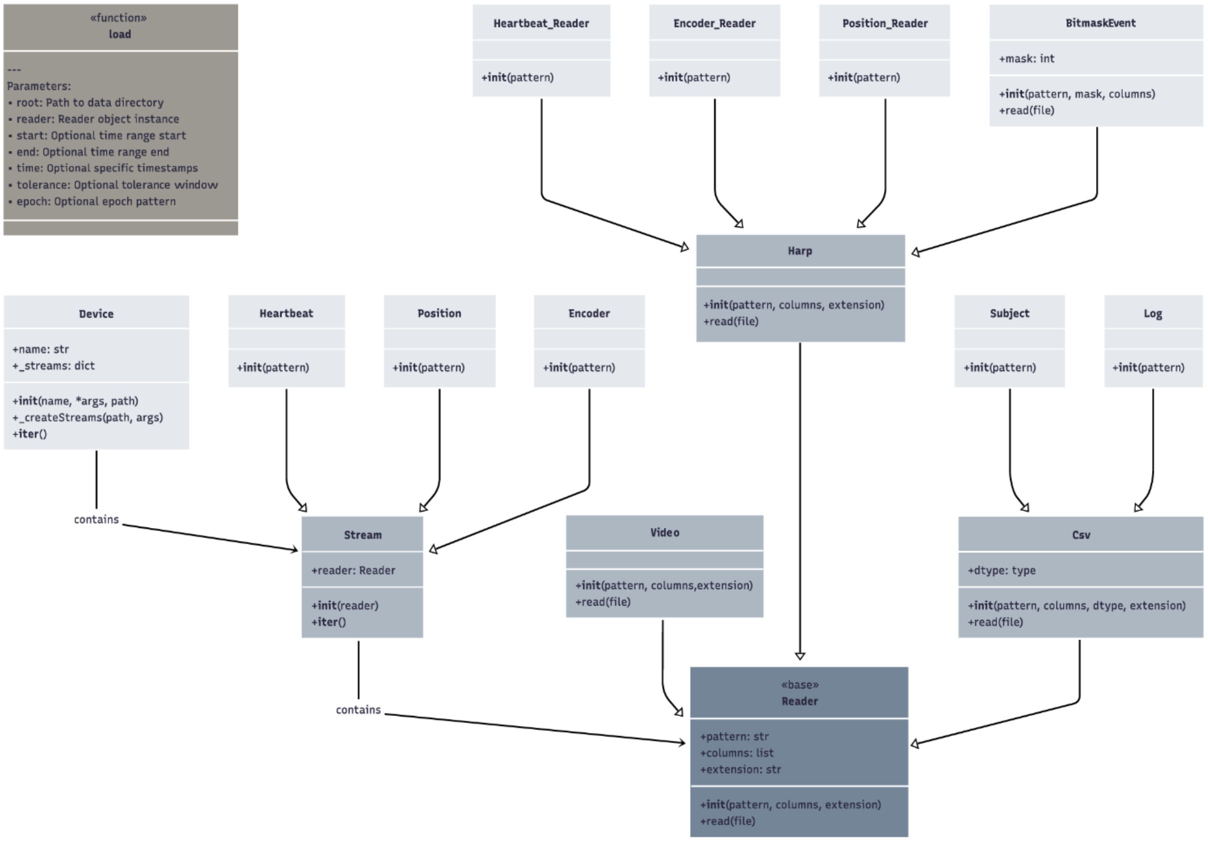


**Extended Data Figure 2 – Aeon low-level Python API**

A simplified, partial UML diagram representation of the Aeon low-level Python API. In the top-left is the API's primary entry point: the 'load' function. From a user's perspective, accessing the raw data starts with a call to 'load'. 'load' provides a single unified interface to raw data regardless of the underlying data stream source. To retrieve data, a user simply specifies a root data path (which can be a path to a specific acquisition epoch within an experiment, an entire experiment, or multiple entire experiments), a 'Reader' object associated with the data to be retrieved, and the desired time range (a start and end time, or a set of specific timestamps with some tolerance).

From the bottom, we illustrate key classes in the API, with subclass relationships indicated by arrows pointing from children to parent classes. As mentioned, each data source is called a 'Stream' and has an associated 'Reader'. For convenience, 'Streams' can be grouped into 'Devices' -- e.g. a 'Patch Device' has an associated 'Feeder Stream' for pellet delivery and 'Encoder Stream' that measures patch wheel rotation. Also shown here are the 'Heartbeat Stream', which contains the clock signal that gets passed to every single data source we record from to ensure sub-millisecond time-alignment for all of our data, and the 'Position Stream', which is a software stream that records online SLEAP pose + identity model inference, containing full ID + pose estimation for each subject in an experiment. Corresponding 'Reader' subclasses are shown alongside these Streams. The purpose of introducing the 'Stream' abstraction, rather than relying solely on 'Reader' classes, is to enable the construction of 'experiment schemas' -- Python objects that contain all data sources for a given experiment (not shown here).

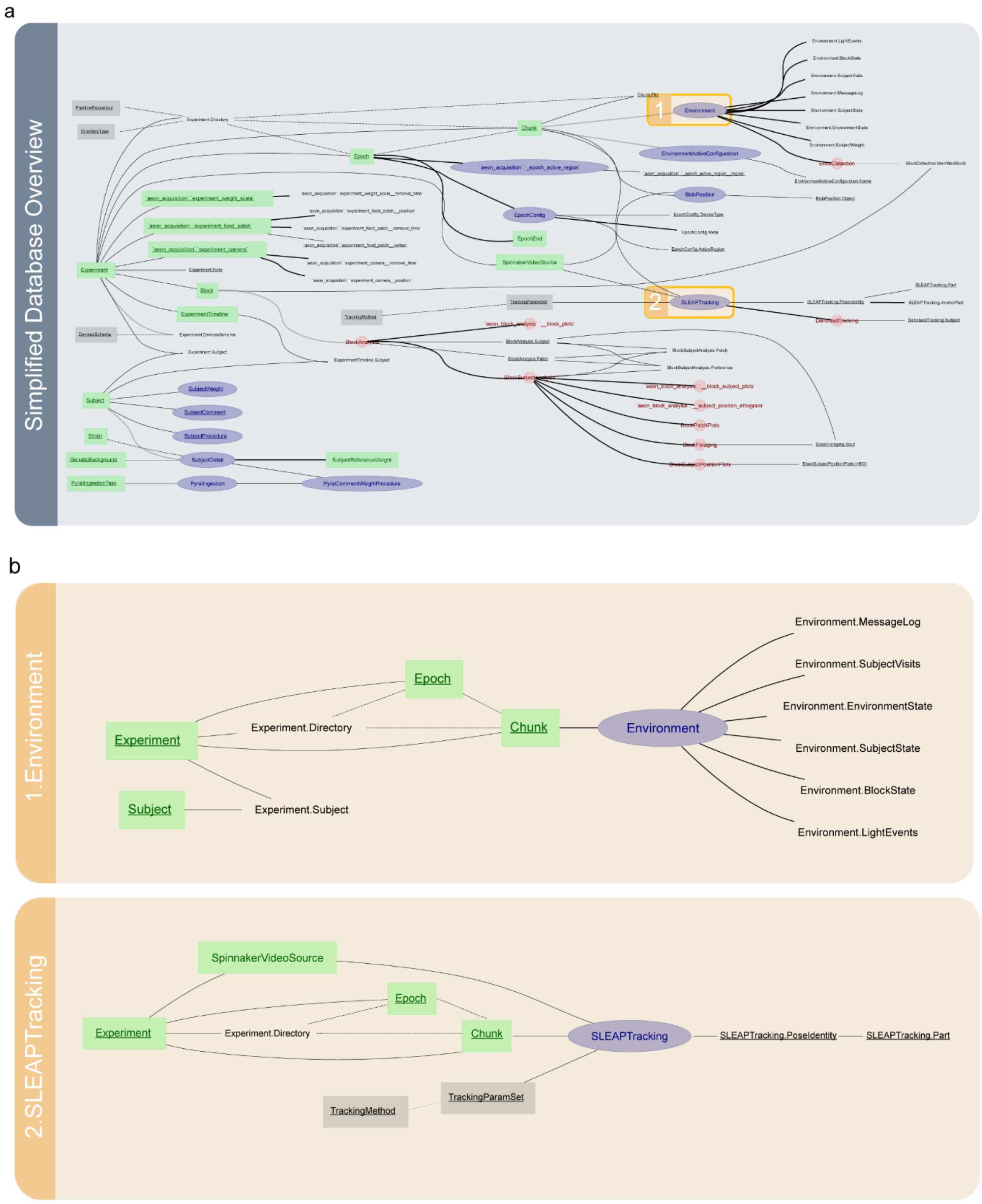


**Extended Data Figure 3 – Architecture of an Aeon DataJoint database**

A simplified, partial ERD illustrating an overview of an Aeon Datajoint database, highlighting particular key tables. A: Zoomed-out overview showing a partial representation of four key schemas in an Aeon datajoint database. B: Zoomed-in view highlighting the ‘Environment’ and ‘SLEAPTracking’ tables, their dependent child tables, which contain processed data derived from specific streams, and the link back to their respective parent source schemas (‘Experiment’ and ‘Subject’).

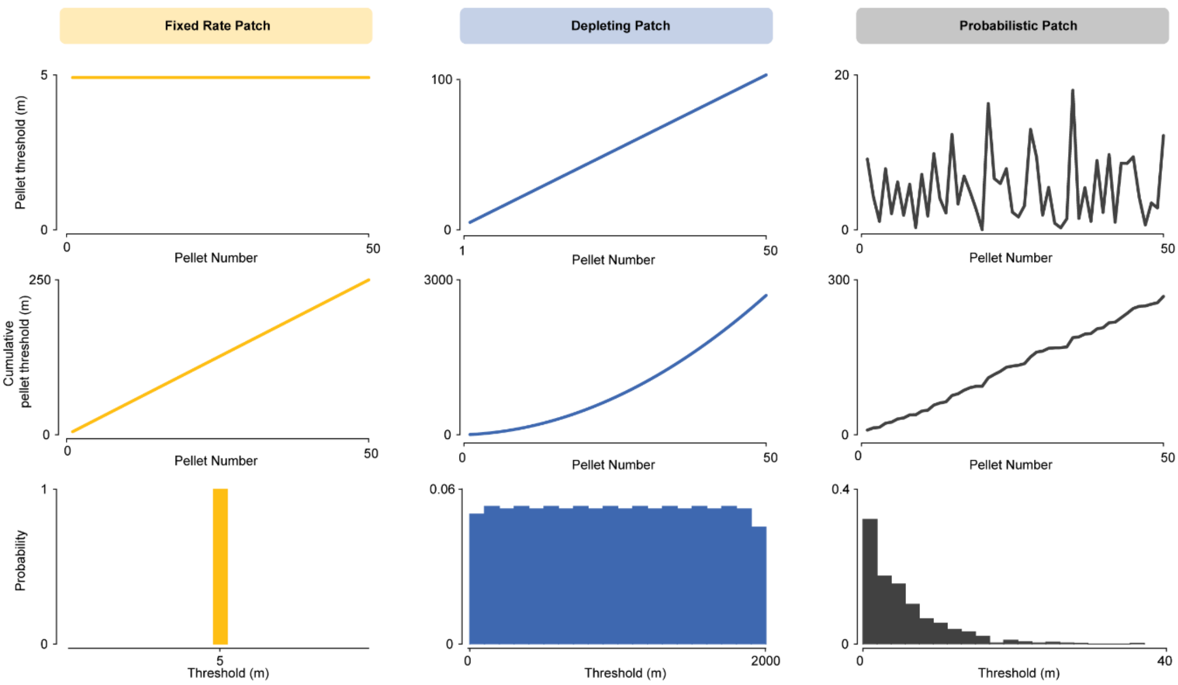


**Extended Data Figure 4 - Example of three foraging patch types implemented in Aeon.**

Fixed Rate Patch: the distance spun between current and next pellet is fixed. In the example it is equal to 5m.

Depleting Patch: the distance between two consecutive pellets starts with a certain value Th0 and increases at every every pellet delivery of a preset delta value. In the example $Th_0$ = *5m* and *delta* = *2m*.

Probabilistic Patch: the distance between two consecutive pellets is drawn from an exponential distribution with set mean. In the example it is equal to 5m.

**Top**: Pellet threshold values for 50 pellet deliveries

**Middle**: Cumulative pellet threshold values for 50 pellet deliveries,

**Bottom**: Estimation of the distribution from where pellet thresholds are drawn, using data from 1000 pellet deliveries.

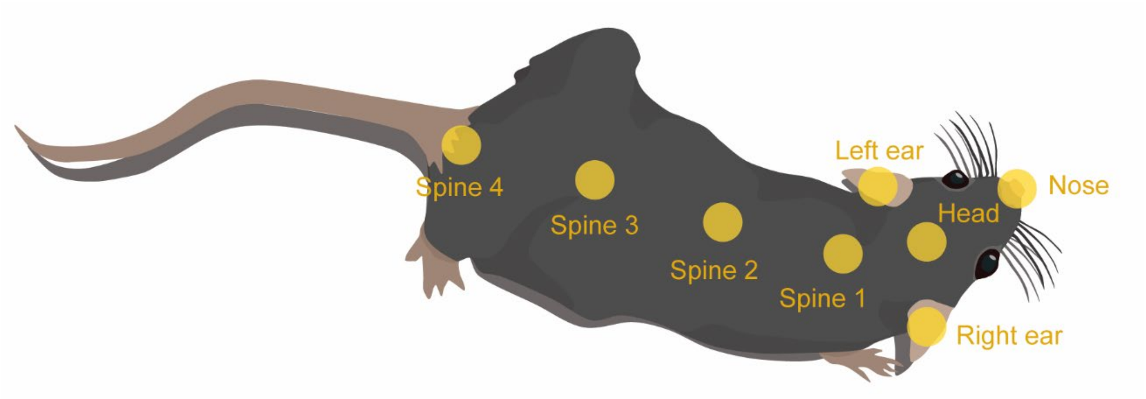


**Extended Data Figure 5 – SLEAP model**

Body parts estimated by the pose SLEAP model used in Fig. 4, 5 and 6.

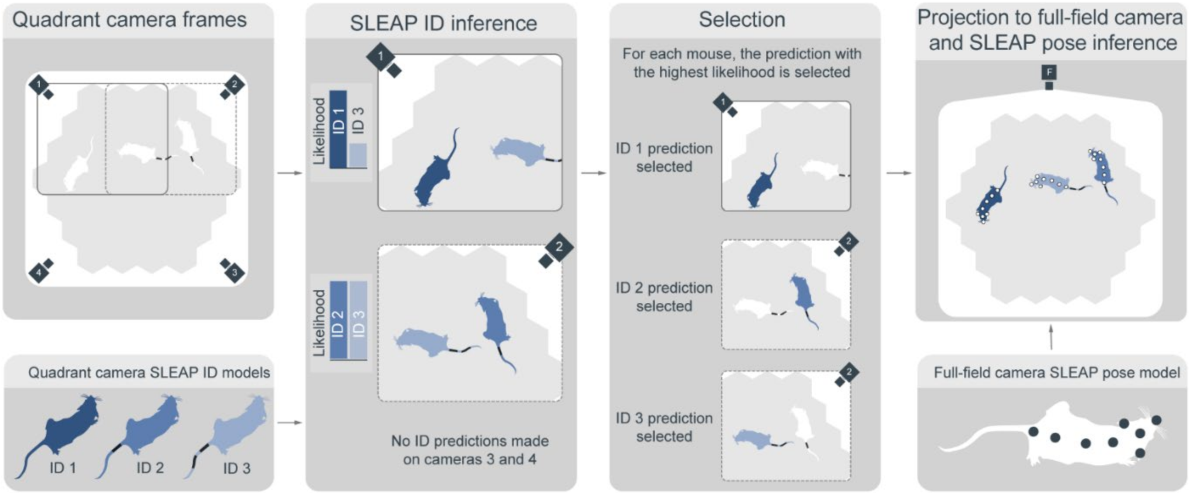


**Extended Data Figure 6 – Schematic of SLEAP-based multi-animal identity and pose tracking.**

(Left) Quadrant cameras capture high-resolution, zoomed-in views of the mice in the arena. (Middle left) A SLEAP identity model runs on all quadrant views, assigning an identity and a likelihood score for each detected mouse. (Middle right) For each mouse, the prediction with the highest likelihood is selected. (Right) These selected identities are then projected back onto the full-field camera view, where SLEAP's pose-inference model estimates multi-point skeletons for each mouse. The final output combines both pose and identity assignments.

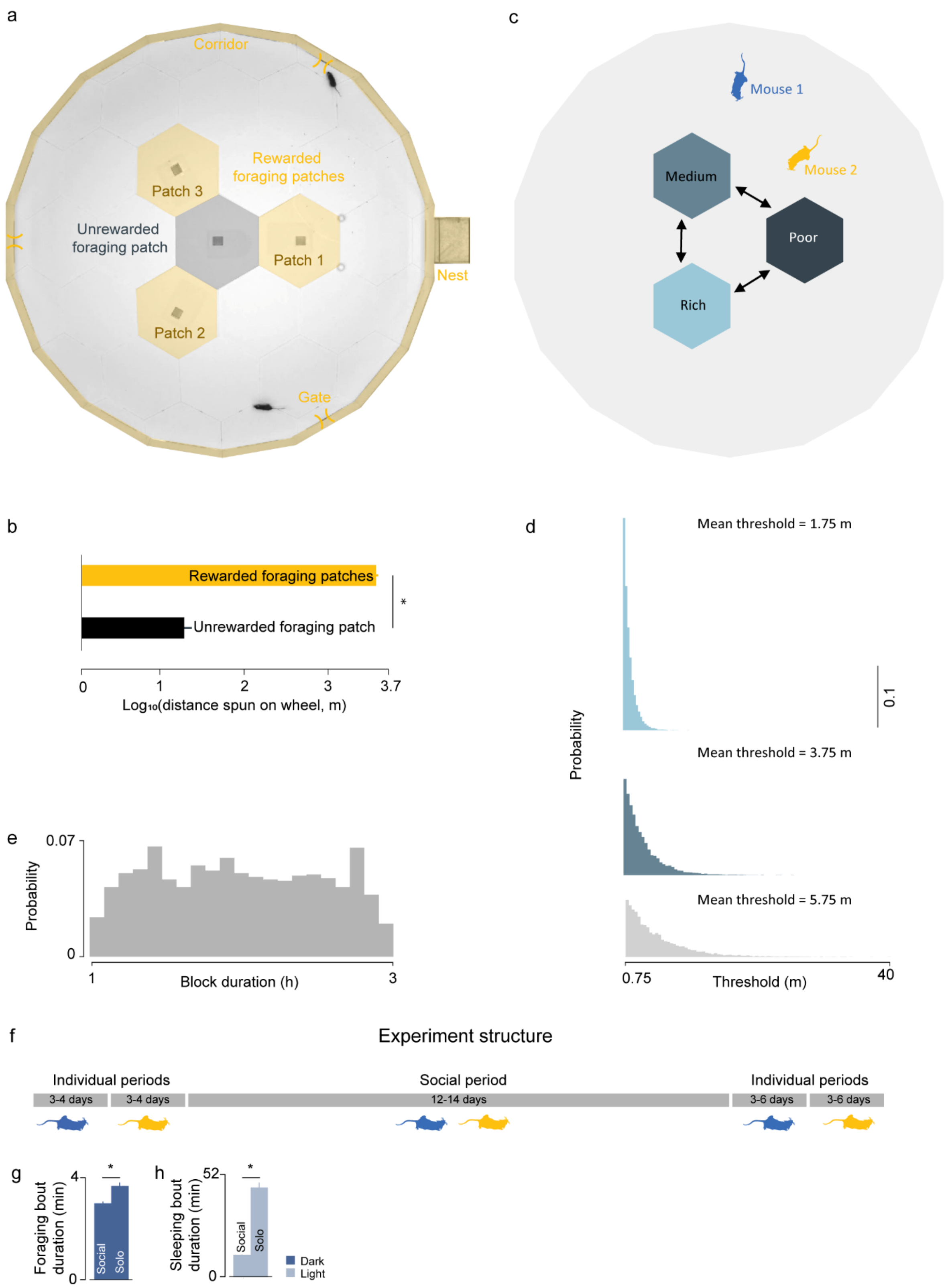


**Extended Data Figure 7 - Social foraging assay**

**a.** Habitat configuration used for the experiments described in Fig. 6d,e. Most experiments were performed with three foraging patches. In a subset of the experiments a fourth patch with an inactive pellet dispenser unit was added as a control. The number of gates could vary between 1or 3.

**b.** Comparison of the distance spun on foraging wheel located on a patch with the dispenser active or inactive. Mice spun a substantially longer distance on patches in which the pellet dispenser unit was active (t-test p-value = 0.0004)

**c.** Schematic of the assay logic of the experiments reported in Fig. 6d,e. Mice were housed in an arena with three foraging patches with different mean pellet threshold: easy patch (mean threshold = 1.75 m), medium patch (mean threshold = 3.75 m) and hard patch (mean threshold = 5.75m). Instantaneous pellet thresholds were sampled from the exponential distributions in panel d. Every ~ 1 to 3 hours (see block durations distribution in panel e), patch difficulty changed.

**d.** Histograms of the distributions of pellet distance thresholds used in the experiments reported in Fig. 6d,e.

**e.** Distribution of block duration of the experiments reported in Fig. 6de.

**f.** Timeline of the experiments reported in Fig. 6d,e.

**g.** Mean foraging bout duration bouts durations during solo and social periods. (* $p<0.05$).

**h.** Mean sleeping bouts durations during solo and social periods. (* $p<0.05$).

Error bars: standard error of the mean

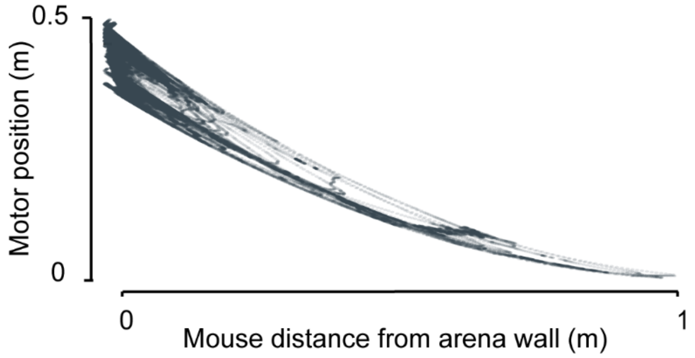


**Extended Data Figure 8**

Plot showing the strong anticorrelation between motor position and mouse distance from the wall (Pearson Correlation Coefficient = 0.93, p-value = 0)

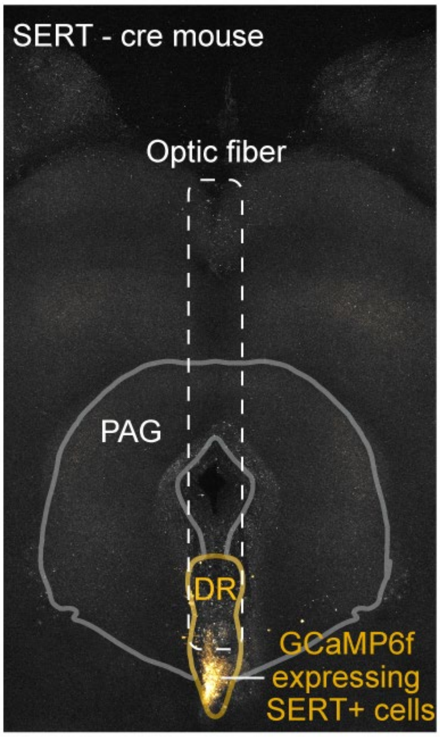


**Extended Data Figure 9 – Fiber photometry optic fiber placement**

Coronal brain section showing optic fibre and viral injection locations for the experiment in Fig. 7d. PAG: Periaqueductal gray. DR: Dorsal Raphe.

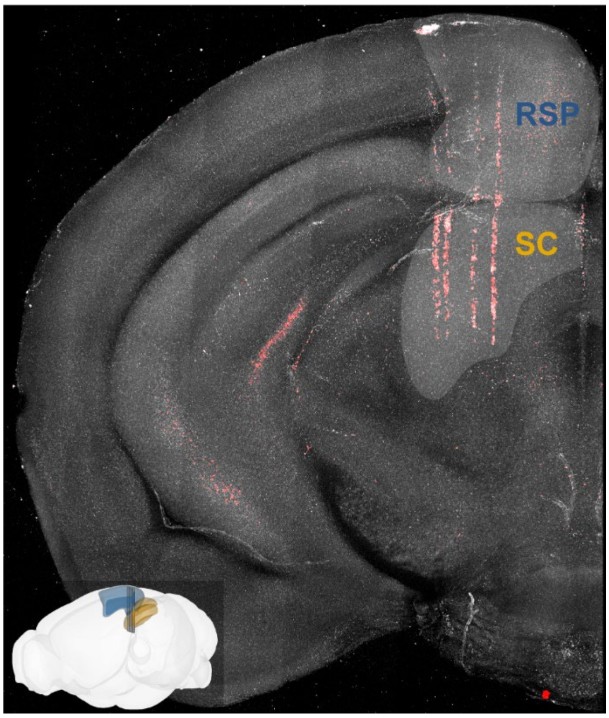


**Extended Data Figure 10**

Histological reconstruction and schematic of probe location (inset) for the electrophysiological recordings shown in Fig. 8.

#### EXTENDED DATA VIDEOS

##### Extended Data Video 1 – Mouse weight measurement

Example of weight measurement event from the scale during a mouse visit to the habitat nest modules. Blue: stable readings. Grey: unstable readings.

##### Extended Data Video 2 – Nesting behaviour

Video showing a mouse engaged in nest building behaviour inside the nest module. Playback speed: 70x.

##### Extended Data Video 3 – Habitat Exploration

Footage of a mouse exploring an Aeon habitat and interacting with the foraging patch modules during a two-patch foraging assay.

##### Extended Data Video 4 – Escape behaviour

Example of a sound-evoked escape episode in an Aeon habitat.

##### Extended Data Video 5 – Drinking behaviour

Example of a drinking episode from the waterspout located inside the nest module.

##### Extended Data Video 6 – Foraging behaviour

20 distinct foraging events from four mice during the interaction with the foraging wheel. All videos are time-aligned to the pellet delivery time.

##### Extended Data Video 7 – Example of digging to threshold foraging event

Left: Example of a mouse spinning the foraging patch module wheel during the digging to threshold foraging assay. Right: wheel position measured by the motion sensor coupled to the wheel. Aqua dot represents pellet delivery time.

##### Extended Data Video 8 – Two patch foraging assay: days one to three

Each panels represent one of the first three days of a two-patch foraging assay. Top. Mouse trajectory: gold; mouse position: red. Middle. Light grey: foraging patch one; dark grey: foraging patch 2. Colourbar indicate the daylight phase in the room: grey represents light phase and blue dark phase.  Playback speed: 1920x.

**Extended Data Video 9 – Examples of HMM states**

Examples of HMM states extracted using the method illustrated in Fig. 5. For each state, the model could reliably identify multiple instances that map on interpretable behavioural syllable.

**Extended Data Video 10 – Multi-animal identity and pose tracking**

Exploration period of two mice performing a three-patch foraging assay. Body parts tracked using SLEAP have been overlayed to each mouse. Colours (gold or blue) indicate the mouse tracked identity.

**Extended Data Video 11 – Aggression bout**

Example of an aggression bout between two mice in an Aeon habitat.

**Extended Data Video 12 – Social encounter and retreating behaviour**

Example of an instance of social encounter and retreating event in the corridor module, used to identify the dominant and the subordinate mouse.

**Extended Data Video 13 – Neuropixels commutation-translation module**

Video illustrating the functioning of the Neuropixels commutation translation module. Top left: tethered mouse chronically implanted with a Neuropixels 2.0 probe while exploring an Aeon habitat. The colour of the trajectory indicates the position of the commutator along the linear rail as illustrated in the top-right panel. Bottom: Position of the commutator along the linear rail and mouse heading direction estimated by the headstage IMU sensor.

**Extended Data Video 14 – Neuropixels recording during week-long foraging assay**

Raw voltage traces recorded from the 384 recording sites of a Neuropixels 2.0 probe (**left**) chronically implanted in a mouse performing a foraging assay (**bottom right**). **Top righ**t: zoomed in view of the raw traces highlighted in blue in the left-hand panel.
